# NLRP1 inflammasome activation in skin equivalents reveals mechanistic insights into the roles of keratinocytes in psoriasis

**DOI:** 10.1101/2025.06.25.661501

**Authors:** Michela Di Filippo, Tugay Karakaya, Phil F. Cheng, Paulina Hennig, Marta Slaufova, Petra Boukamp, Steve Pascolo, Julia-Tatjana Maul, Isabel Kolm, Mitchell P. Levesque, Thomas Kündig, Hans-Dietmar Beer

## Abstract

Psoriasis is a major inflammatory skin disease for which a causal therapy is still not available. The pro-inflammatory cytokines interleukin(IL)-1β and IL-36γ are key drivers of the disease phenotype, but the mechanisms underlying their regulation in psoriasis remain poorly understood. Generation of IL-1β activity is regulated by protein complexes, termed inflammasomes. We activated the NLRP1 inflammasome in human keratinocytes cultivated in three-dimensional skin equivalents. NLRP1 activation induced histological and molecular features that are highly reminiscent of psoriasis. Mechanistically, the phenotype was dependent on IL-1, which triggered a pro-inflammatory epidermal-dermal crosstalk. This included induction of expression of IL-36γ, which, together with IL-1β, was released from keratinocytes through NLRP1-induced gasdermin D pores. The *in vivo* relevance of these findings is reflected by the expression of the NLRP1 sensor and signs of inflammasome activation in lesional skin of psoriatic patients. Finally, we discovered endogenous cytoplasmic double stranded (ds) RNA, recently associated with cellular perturbations in psoriasis, as a novel activator of the NLRP1 inflammasome in human keratinocytes.

Our results identify a novel endogenous double-stranded RNA-mediated NLRP1-IL-1-IL-36γ signaling axis relevant in psoriasis and suggest targeting of this pathway as a promising treatment strategy.

## INTRODUCTION

Psoriasis is a chronic, incurable inflammatory disease affecting 2-3% of the global population and characterized by sharply demarcated erythematous and scaly skin lesions [1, 2]. While psoriasis primarily affects the skin, it is increasingly recognized as a systemic disorder [3]. At the histological level, skin lesions are characterized by T cell infiltration and a massive thickening of the epidermis due to increased proliferation and aberrant differentiation of keratinocytes, the main cell type of the epidermis. Psoriasis is a highly complex genetic disorder, with more than 80 associated genomic risk loci identified, and presents in various clinical form [3, 4]. Psoriasis vulgaris (PV) is the most common type of psoriasis, with IL-23 and IL-17 pathways playing key pathogenic roles and being effectively targeted [2]. Generalized pustular psoriasis (GPP) is rare but live-threatening, frequently caused by loss-of-function mutations of IL-36 receptor antagonist (IL-36RA) [5]. IL-36RA inhibits the IL-36 receptor (IL-36R) that is activated by the pro-inflammatory IL-1 family members IL-36α, -β, and -γ. Consequently, a humanized blocking antibody for IL-36R was recently approved for the treatment of GPP [6, 7]. Interestingly, keratinocytes are the primary cells expressing IL-36R and, through their production of IL-36γ, serve as the main source of IL-36 activity in psoriatic lesions [7]. IL-36γ is initially expressed as a biologically inactive pro-protein, which is activated by the cysteine protease cathepsin S in keratinocytes [8]. However, IL-36γ lacks a signal peptide for secretion by the canonical ER/Golgi-dependent pathway and release of the cytokine by keratinocytes remains poorly understood.

The potent pro-inflammatory cytokine IL-1β is the prototypic IL-1 family member [9]. While IL-1β is a key inducer of inflammatory responses required for protection from pathogens and repair processes, its excessive or chronic activity contributes to numerous inflammatory conditions [9]. IL-1β exerts its biological effects by binding to IL-1 receptor type I (IL-1R1), which is ubiquitously expressed. Individuals with over-activation of IL-1R1, resulting from loss-of-function mutations of *IL1RN* encoding the IL-1 receptor antagonist (IL-1RA), suffer from various symptoms, including pustular skin eruptions resembling pustular psoriasis [10, 11]. Furthermore, single nucleotide polymorphisms (SNPs) of *IL1RN* are associated with an increased susceptibility to psoriasis [12]. Although these genetic and other experimental data suggest a critical role of IL-1 signaling in psoriasis, the underlying molecular and cellular mechanisms are incompletely understood [13–15].

IL-1β exerts its proinflammatory effects following proteolytic activation by the cysteine protease caspase-1, which is activated upon inflammasomes assembly [16]. Inflammasomes are a group of protein complexes that consist of caspase-1, the adaptor protein apoptosis-associated speck-like protein containing a caspase recruitment domain (ASC), and a sensor protein, such as nucleotide-binding domain, leucine-rich-containing family, pyrin domain-containing-1 (NLRP1), NLRP3, or absent in melanoma 2 (AIM2). These sensors act as pattern recognition receptor (PRR), detecting stress signals like pathogen-associated molecular patters (PAMPs), damage-associated molecular patters (DAMPs), or homeostasis-altering molecular processes (HAMPs) [17]. Sensing of the stressor induces oligomerization of the inflammasome receptor, leading to the recruitment of ASC and its assembly into large aggregates known as ASC specks, a hallmark of inflammasome activation. This process is followed by binding of caspase-1 and its proximity-induced self-activation [16]. Then, caspase-1 cleaves and activates proIL-1β and proIL-18, as well as gasdermin D (GSDMD). The latter forms pores in the plasma membrane, which are required for the release of active IL-1β and IL-18. Like IL-36γ, IL-1β and IL-18 lack a signal peptide for secretion by the canonical pathway. Furthermore, GSDMD pores can induce a lytic type of cell death, termed pyroptosis, which supports inflammation [18]. Inflammasomes play crucial roles in immune defense, but they also contribute to common inflammatory conditions [19].

NLRP1 is considered the main inflammasome sensor in human skin and keratinocytes [20]. SNPs of *NLRP1* are associated with several inflammatory diseases that primarily affect the skin, such as vitiligo or psoriasis [21].

NLRP1 is activated downstream of the ribotoxic stress response (RSR) [22, 23]. Different toxins, antibiotics, as well as UVB radiation, which was identified as the first stimulus for NLRP1 activation, activate the RSR [24, 25]. Furthermore, NLRP1 is activated upon viral infection, induced by either proteolytic cleavage in the amino-terminal fragment through viral 3C proteases or direct binding of long viral double-stranded (ds) RNA [26, 27]. In contrast, NLRP1 activation is restrained or even completely inhibited by binding to oxidized thioredoxin or dipeptidyl peptidases (DPP) 8/9, respectively [28, 29]. Consequently, inhibitors of DPP8/9, which target the active site where NLRP1 is bound, induce NLRP1 inflammasome activation [28].

The NLRP1 pathway is only poorly conserved in the skin of mice [30, 31]. To investigate NLRP1 activation in a physiologically relevant context, we pharmacologically activated NLRP1 in human primary keratinocytes (HPKs) cultivated in a 3D organotypic model of human skin, known as skin equivalent (SE) [32]. We observed that NLRP1 activation in SEs induced an IL-1-driven histological and molecular phenotype sharing key characteristics with psoriasis. We identified IL-36γ as a cytokine regulated downstream of NLRP1 and secreted through GSDMD pores. Additionally, we identified short endogenous dsRNA, a stress factor in psoriasis, as a novel NLRP1 activator. Together, these findings highlight a critical role of NLRP1 and keratinocytes in the pathogenesis of psoriasis and suggest that SEs with activated NLRP1 provide a novel model for investigating the inflammatory skin disease and for exploring new therapeutic approaches.

## RESULTS

### SEs with CRISPR/Cas9-modified HPKs represent a physiological *in vitro* skin model

Unlike human keratinocytes, murine keratinocytes in culture do neither express significant levels of NLRP1 nor of IL-1β [30]. Furthermore, the molecular mechanisms regulating activation of NLRP1 are only partially conserved between humans and rodents [31]. Organotypic skin culture based on human primary fibroblasts (HPFs) and keratinocytes (HPKs) provides a physiological model for human skin, particularly concerning the morphology of the epidermis [33]. More recently, a fibroblast-derived matrix-based full-thickness SE has been developed, closely mimicking the histology of human skin with epidermal stratification and keratinocyte differentiation, offering exceptional stability and reproducibility [32, 34]. We cultured polyclonal CRISPR/Cas9 control knockout HPKs in these SEs and performed RNA sequencing after enzymatic separation of the dermis and epidermis. To evaluate the physiological relevance of these SEs for studying human skin, we compared the RNA sequencing data with that from human skin and observed a similar expression pattern (Fig. 1A) [35]. Furthermore, histological analysis of the expression of epidermal differentiation markers and comparison with human skin revealed that scaffold-free SEs with CRISPR/Cas9 control knockout HPKs represent a physiological model for human skin (Fig. 1B).

**Fig. 1:**
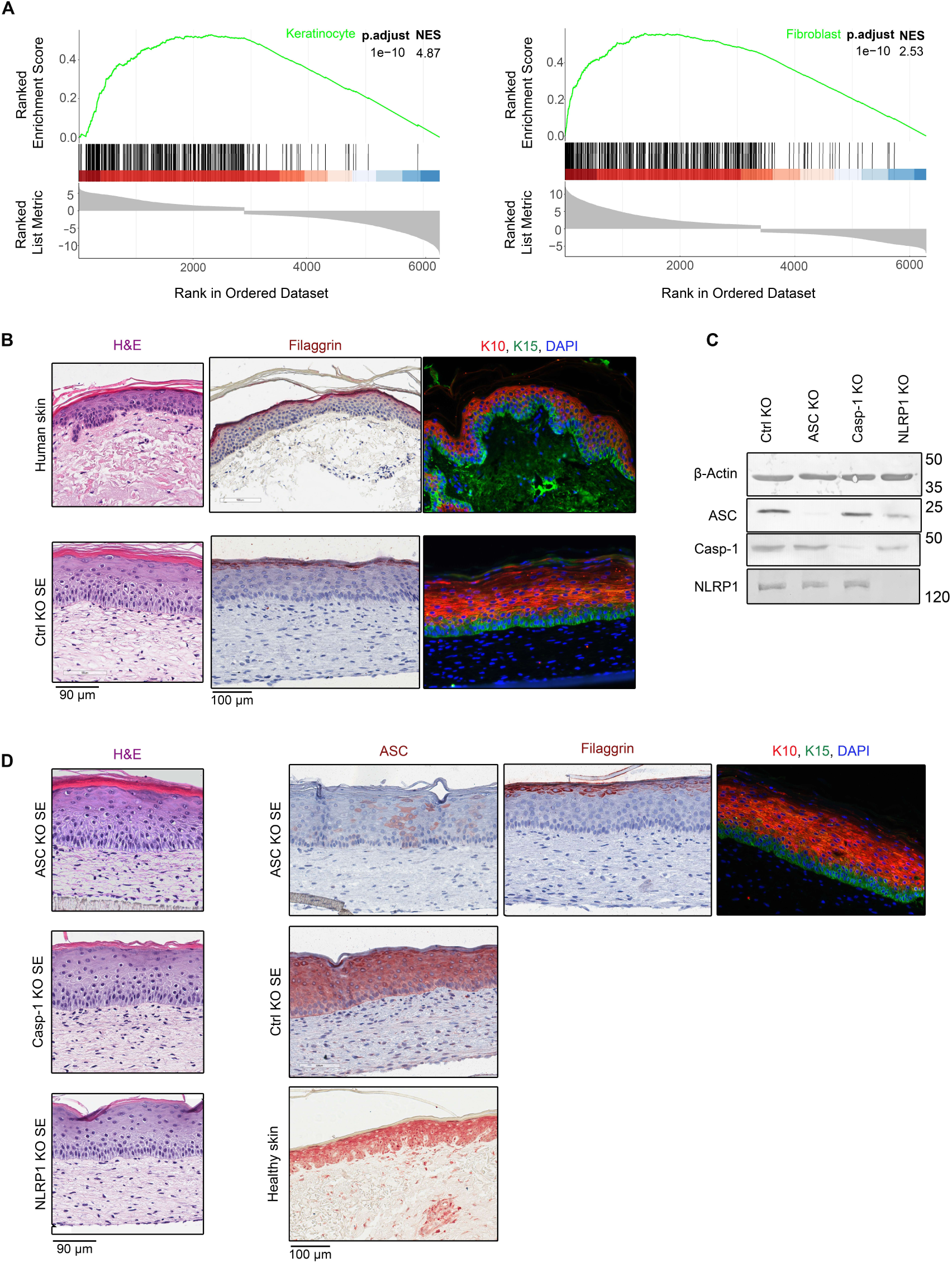
Skin equivalents (SEs) with CRISPR/Cas9-modified HPKs resemble human skin. SEs were generated with CRISPR/Cas9-modified HPKs. A control non-targeting sgRNA was used (Ctrl KO) or sgRNAs targeting the *ASC*, *caspase-1*, or *NLRP1* gene. **(A)** GSEA graph demonstrates that gene expression of control HPKs (left) or HDFs (right) in SEs resembles the one in the epidermis or dermis in human skin. Genes were ordered according to the differential expression in HPKs over HDFs for the epidermis and vice versa for the dermis of the SEs. Gene sets “keratinocytes” and “fibroblasts” were obtained from [35]. Genes in the red area represent positive NES indicating upregulation of the gene set in the ranked list, highlighting its enrichment. **(B,C,D)** CRISPR/Cas9-modified control and NLRP1 inflammasome knockout HPKs form an epidermis in SEs. **(B)** H&E, immunohistochemical or -fluorescence staining of SEs with control knockout HPKs and of healthy human skin for the expression filaggrin and keratin10 and −15 (K10, K15). DAPI stains nuclei. Scale bar 90 µm or 100 µm. **(C)** Western blot for expression of the indicated proteins by polyclonal knockout HPKs. **(D)** (Left) H&E staining of SEs with ASC, caspase-1 and NLRP1 knockout HPKs. Scale bar 90 µm. (Right) Immunohistochemical or -fluorescence staining of SEs with ASC knockout HPKs for expression of ASC, filaggrin and keratin10 and −15 (K10, K15). SEs with Ctrl knockout HPKs and healthy skin are used as controls for ASC staining. DAPI stains nuclei. Scale bar 100 µm. **(B,C,D)** Data are representative of at least 3 independent experiments. GSEA: gene set enrichment analysis; H&E: hematoxylin and eosin; HDFs: human dermal fibroblasts; HPKs: human primary keratinocytes; KO: knockout; NES: normalized enrichment score; SE: skin equivalent.

Then, we also generated polyclonal CRISPR/Cas9 knockout HPKs lacking expression of NLRP1, ASC, or caspase-1 (Fig. 1C,D) and demonstrated that these knockout HPKs also formed a stratified epidermis in SEs (Fig. 1D).

### Talabostat treatment induces NLRP1 activation in suprabasal keratinocytes

In unstimulated keratinocytes, NLRP1 is inhibited by binding to dipeptidyl peptidase (DPP) 8 and 9 [36]. Talabostat is a DPP8/9 inhibitor and releases NLRP1 from the peptidase in a specific manner by binding DPP8/9’s active site [28]. Treatment of HPKs with talabostat (0.3 µM) induced secretion of high levels of IL-1β dependent on the expression of the NLRP1 inflammasome components NLRP1, ASC, and caspase-1 (Fig. 2A). In a similar manner, talabostat (0.3 µM) induced IL-1β secretion from control HPKs in SEs, which was absent in dermal equivalents, and strongly reduced in SEs with NLRP1 inflammasome knockout HPKs (Fig. 2B). As the ASC knockout most efficiently abolished IL-1β secretion, we used ASC knockout cells for all the subsequent experiments with SEs. We stained talabostat- and mock-treated control and ASC knockout SEs for ASC specks, a second read-out for inflammasome activation. Surprisingly, we detected ASC specks exclusively in suprabasal HPKs upon NLRP1 activation induced by talabostat (Fig. 2C). In 2D monoculture, the NLRP1 inflammasome can be efficiently activated in proliferating HPKs without priming [24, 28], suggesting that in 3D culture this would occur in basal cells. However, also in *ex vivo* human skin, we detected talabostat-induced ASC speck formation in suprabasal cells (Fig. 2D), demonstrating that SEs are a better model for NLRP1 activation in human epidermis than HPKs in 2D monoculture.

**Fig. 2:**
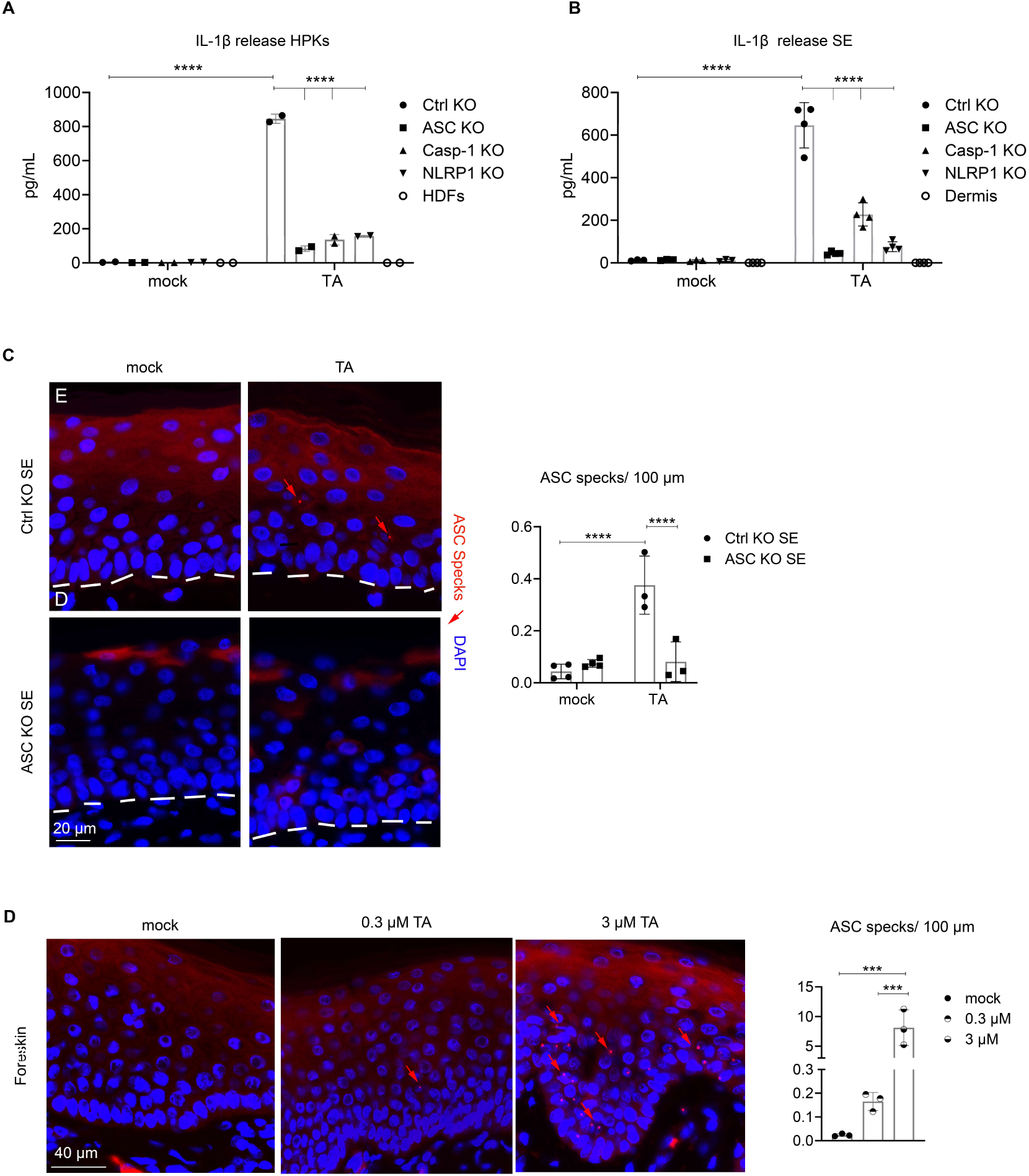
Talabostat activates the NLRP1 inflammasome in suprabasal keratinocytes. Control and knockout HPKs in **(A)** monoculture or **(B)** SEs were treated with 0.3 µM talabostat for 3 days and analyzed for secretion of IL-1β by ELISA. **(A)** HDFs or **(B)** dermal equivalents (dermis) served as controls. **(C)** ASC immunofluorescence staining of SEs with control or ASC knockout HPKs 3 days after mock- or treatment with 0.3 µM talabostat. ASC specks (red arrow) were quantified. Scale bar 20 µm. **(D)** ASC immunofluorescence staining of healthy human skin mock-treated for 3 days *ex vivo* or with the indicated concentrations of talabostat. The number of ASC specks was quantified. Scale bar 40 µm. Data are representative of **(A)** 2 or **(B,C,D)** at least 3 independent experiments. P values were calculated with **(A,B,C)** two-way ANOVA and **(D)** one-way ANOVA. **(A)** N=2, **(B,C,D)** N≥3, (∗∗∗∗P < 0.0001, ∗∗∗P ≤ 0.001, ∗∗P ≤ 0.01, and ∗P ≤ 0.05, ns = not significant). TA: talabostat

Then, to confirm the specificity of talabostat as an NLRP1 activator in SEs, we treated the model with a high concentration of talabostat (10 µM) for 3 days, replenishing the medium with fresh talabostat daily. Although this treatment caused complete epidermal atrophy in SEs with control HPKs, those with ASC knockout cells were protected (Fig. S1A). In contrast, NLRP1 activation in keratinocytes via the RSR or transfection of the synthetic dsRNA poly(I:C) also activates other cell death pathways [22, 27].

In conclusion, these experiments demonstrate that talabostat is a very specific stimulus for NLRP1 activation in HPKs cultivated in SEs and therefore, that this system represents a physiologically relevant model to address the roles of the NLRP1 inflammasome in human skin.

### NLRP1 activation in SEs induces expression of pro-inflammatory genes, particularly in the dermis

To investigate the consequences of NLRP1 activation, SEs were either mock-treated or with the NLRP1 activator talabostat (0.3 µM) for 3 days in triplicate. The SEs were prepared with either control or ASC knockout HPKs. Then, we separated dermis and epidermis and analyzed gene expression by next-generation-sequencing (NGS). Talabostat treatment caused a statistically significant regulation of 168 genes in epidermal keratinocytes, in an ASC-dependent manner (Fig. 3). These genes code for proteins which are either involved in or associated with differentiation of keratinocytes, such as transglutaminase 1 (TGM1) and small proline-rich proteins (SPRRs), or are involved in defense or innate immunity, such as defensins, S100 proteins, IL-1β, or IL-36γ (Fig. S1C). Surprisingly, although NLRP1 activation occurred in keratinocytes, talabostat treatment resulted in the significant deregulation of 1048 genes in dermal fibroblasts, in an ASC-dependent manner. These dermal genes are involved in pathways related to immunity, defense, cell activation, and cytokine-induced signaling, including C-C motif ligand (CCL) or C-X-C motif ligand (CXCL) chemokines, matrix metalloproteinases (MMPs), IL-36α and -β, or IL-8. As talabostat activates NLRP1 specifically in keratinocytes, expression of dermal genes is presumably induced by IL-1β or IL-18, which are the known secreted effector proteins of inflammasome activation. This experiment demonstrates that NLRP1 activation in keratinocytes in a physiological skin model without any immune cells induces expression of a pronounced pro-inflammatory signature, particularly in dermal fibroblasts, through paracrine cytokine signaling. Therefore, it seems reasonable that also in human skin, NLRP1 inflammasome activation in keratinocytes might induce an inflammatory response and the infiltration of immune cells, the latter possibly regulated through the pro-inflammatory dermal expression pattern.

**Fig. 3:**
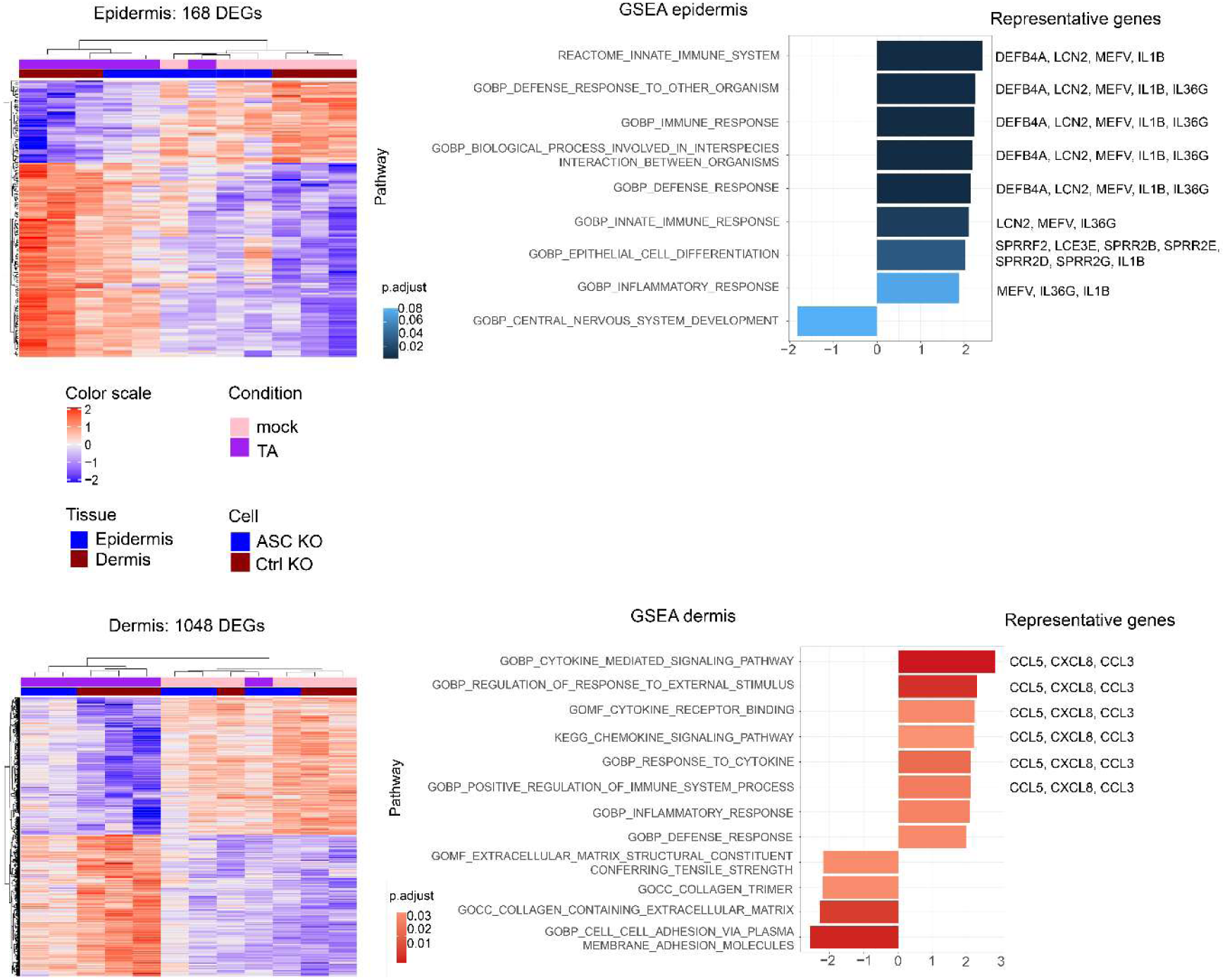
NLRP1 activation in SEs induces a pro-inflammatory signature. SEs with control knockout or ASC knockout HPKs (triplicates) were mock- or talabostat-treated (0.3 μM) for 3 days. Epidermis and dermis were separated and mRNA expression characterized by NGS. (Left) Heatmaps of keratinocytes with 168 differentially expressed genes and of fibroblasts with 1048 genes, upon talabostat-induced ASC-dependent NLRP1 activation, are shown. (Right) GSEA reveals strongly induced and repressed pathways. Representative genes are leading edge genes of these pathways chosen for further analysis. DEGs: differentially expressed genes; GSEA: gene set enrichment analysis.

### IL-1 is the main effector of NLRP1 inflammasome activation and induces IL-36γ expression

To ensure the reproducibility of our RNA-seq findings, which were initially obtained using SEs derived from a single donor, we extended our analysis to multiple donors. We generated SEs with HPKs from 5 different donors, treated them with talabostat as before, and analyzed mRNA levels of representative genes of the regulated pathways. Despite some donor-to donor variability, all genes showed consistent upregulation (Fig. S2), confirming that NLRP1 activation and the induction of the downstream pathways are representative, reproducible, and donor-independent.

To identify the main driver of the observed inflammatory signature, we focused on IL-1β, the key downstream effector of inflammasome activation [37], and addressed its role in the skin model. We treated SEs either with IL-1 (at the maximum concentration of IL-1α and -β secreted following talabostat treatment, Fig. 5A), talabostat alone, or anakinra plus talabostat. Anakinra, a recombinant human IL-1RA, binds to IL-1R1 and prevents signaling by IL-1α and -β. We then analyzed mRNA levels of the previously defined representative genes. As expected, anakinra inhibited talabostat-induced expression of all genes after 3 days (Fig. 4A), and to a lesser extent after 1 day (Fig. S3A), confirming IL-1 as the central effector of the NLRP1 inflammasome in keratinocytes concerning mRNA regulation of the representative genes.

**Fig. 4:**
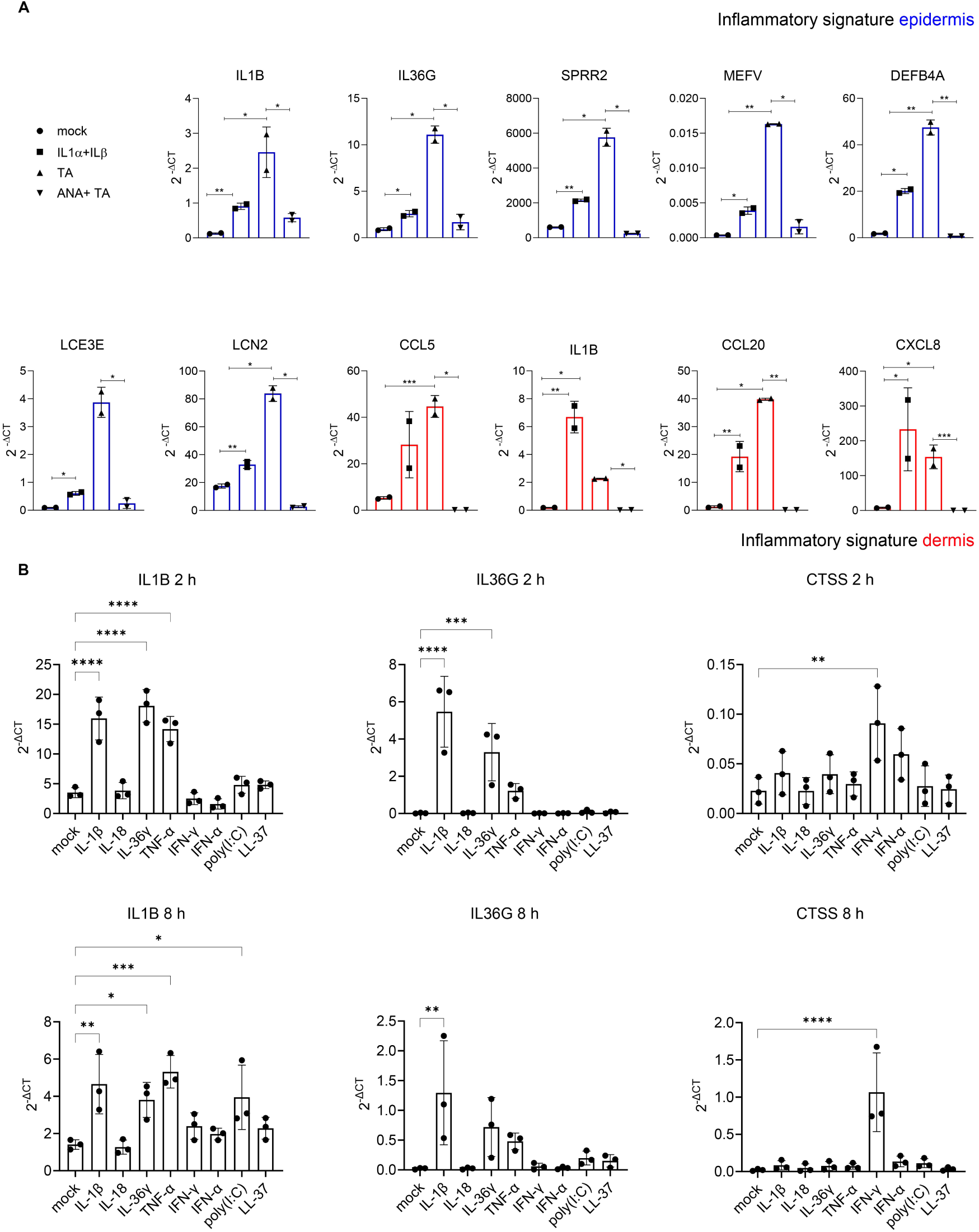
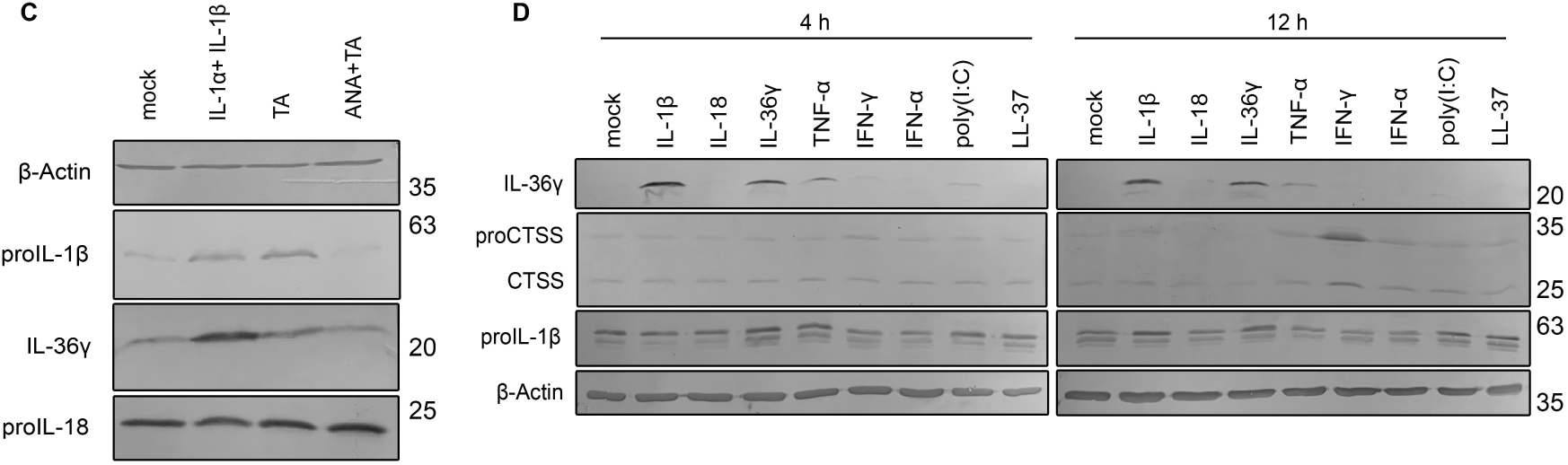
IL-1 is major effector of NLRP1 inflammasome activation. (A,C) SEs were mock-treated or with IL-1α/β (1 ng/ml each), talabostat (0.3 μM), or anakinra (10 μg/ml) plus talabostat, for 3 days. **(A)** Expression of mRNA of representative genes of the epidermis (blue) and of the dermis (red) was determined by qPCR related to HPRT expression. **(C)** Protein expression of IL-1β, IL-36γ and IL-18 was analyzed in the epidermis of SEs 3 days after treatment by western blot. Expression of β-Actin served as control. **(B,D)** Starved HPKs from 3 different donors in monolayer were treated with IL-1β (10 ng/ml), IL-18 (20 ng/ml), IL-36γ (100 ng/ml), TNFα (10 ng/ml), IFN-γ (20 ng/ml), IFN-α (10 ng/ml), poly(I:C) (1 μg/ml), or LL-37 (1 μg/ml). IL-1β, IL-36γ and cathepsin S (CTSS) expression was determined at the **(B)** RNA level 2 h (upper panel) and 8 h (lower panel) after stimulation and **(D)** at the protein level after 4 h and 12 h. **(A)** Data are represented by mean ± SD of 2 replicates and are representative of 2 independent experiments, or **(B)** 3 different donors, or a representative blot out of 2 **(C)** or **(D)** 3 donors is shown. P values were calculated with one-way ANOVA (∗∗∗∗P < 0.0001, ∗∗∗P ≤ 0.001, ∗∗P ≤ 0.01, and ∗P ≤ 0.05, ns = not significant). ANA: anakinra; TA: talabostat.

**Fig. 5:**
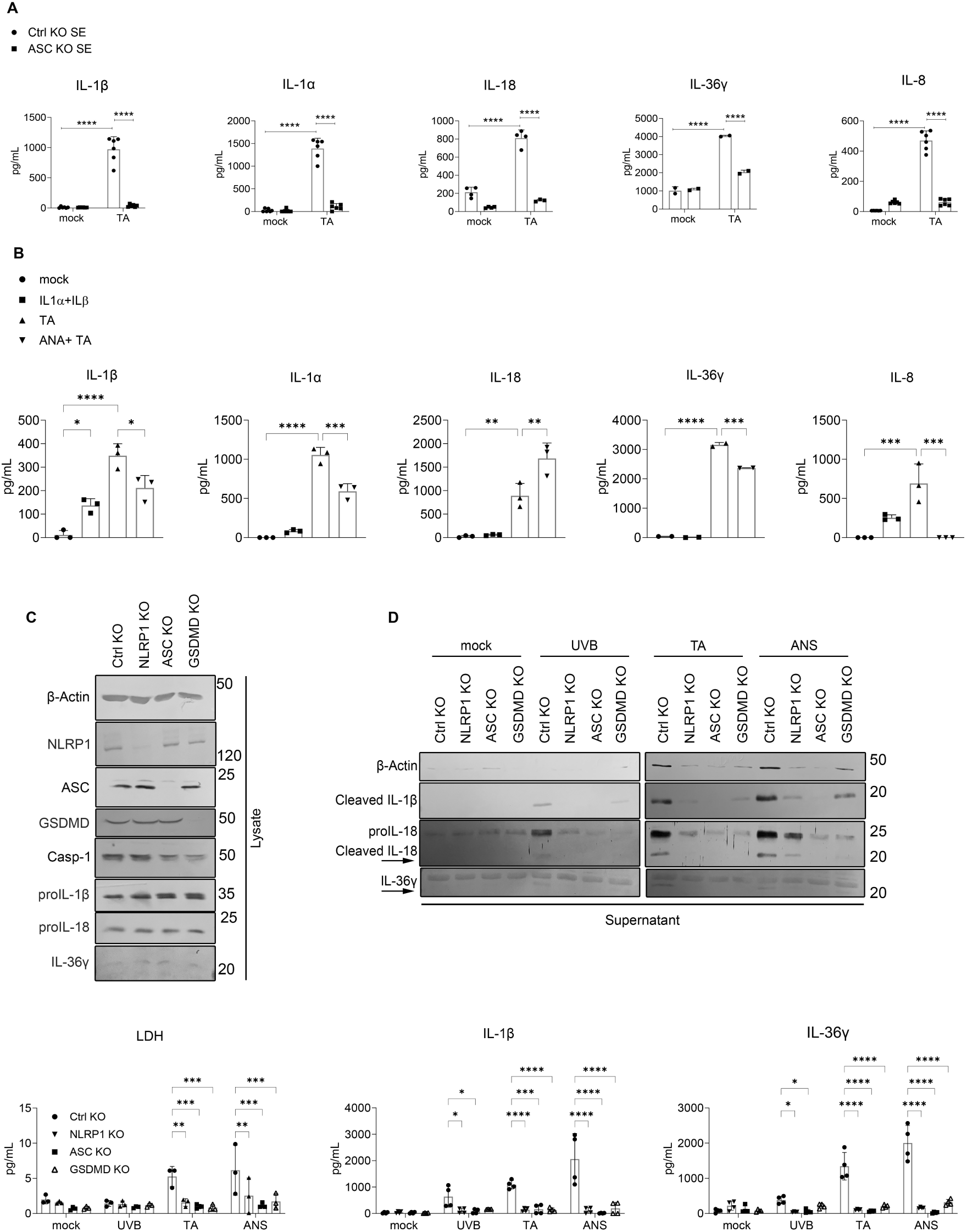
NLRP1 activation regulates IL-36γ release. SEs with **(A)** control or ASC knockout HPKs or **(B)** wild type HPKs were **(A)** mock-treated or with talabostat (0.3 μM), or **(B)** mock-treated or with IL-1α/β (1 ng/ml each), talabostat (0.3 μM), or anakinra (10 μg/ml) plus talabostat, for 3 days. Cytokine release was measured by ELISA. **(C)** Western blot for expression of the indicated proteins by mock-treated polyclonal knockout HPKs in 2D monoculture primed with IL-1α (10 ng/ml) overnight for the induction of IL-36γ expression. **(D)** Knockout HPKs from 3 independent donors in 2D monolayer primed with IL-1α, then mock-treated or irradiated with UVB (86.4 mJ/cm^2^, 6 h), treated with talabostat (10 μM, 12 h), or with anisomycin (1 μM, 5 h) and protein release was characterized by western blot (upper panel) or ELISA (lower panel). Pyroptosis was measured by LDH release (lower panel). **(A,B)** Data are represented by mean ± SD of at least 2 replicates and are representative of 2 independent experiments. **(C,D)** Data are represented by mean ± SD of 3 donors or a representative blot out of 3 is shown. **(A,D)** P values were calculated with two-way ANOVA and **(B)** with one-way ANOVA (∗∗∗∗P < 0.0001, ∗∗∗P ≤ 0.001, ∗∗P ≤ 0.01, and ∗P ≤ 0.05, ns = not significant). TA: talabostat,; ANA: anakinra.

Among the IL-1-regulated genes, we focused on IL-36γ, a pro-inflammatory cytokine with an increasingly recognized role in skin inflammation [38]. In keratinocytes, IL-36γ is cleaved and activated by cathepsin S [8]. To investigate the central roles of IL-1β and IL-36γ, we analyzed regulation of expression of these cytokines, cathepsin S and inflammasome components in starved HPKs in 2D monoculture (cultivated in K-SFM without EGF and BPE), treated with different pro-inflammatory stimuli (Fig. 4B,D and Fig. S3B). Starved HPKs constitutively express NLRP1 inflammasome components, including IL-1β, and IL-1β expression is further increased by IL-1β itself or by IL-36γ. In contrast, IL-36γ is not expressed under basal conditions but is strongly induced at both mRNA and protein level by IL-1β and IL-36γ, demonstrating a crucial role of IL-1β in the regulation of IL-36γ expression (Fig. 4B,D).

In the epidermis of SEs, IL-1 stimulation resulted in the highest IL-36γ protein levels despite relatively low mRNA expression, while talabostat induced strong mRNA upregulation but resulted in only intermediate protein levels (Fig. 4A,C), suggesting that talabostat may influence IL-36γ regulation through additional mechanisms (see below).

### NLRP1 inflammasome activation regulates IL-36γ secretion via GSDMD

To gain a better understanding of IL-36γ regulation in NLRP1-activated SEs, we measured the secretion of IL-36γ, IL-1α, IL-1β, IL-18, and IL-8 at the protein level in the medium of talabostat- or mock-treated SEs (Fig. 5A). As suggested by their mRNA regulation (Fig. 4A), release of all cytokines was induced upon NLRP1 activation in SEs with control HPKs and significantly lower in those with ASC knockout HPKs (Fig. 5A). The analysis of SEs, treated with IL-1, talabostat alone, or anakinra plus talabostat, revealed that IL-8 secretion is induced by IL-1, either directly (by recombinant IL-1) or upon NLRP1 activation, since it is completely suppressed by anakinra (Fig. 5B). Indeed, this aligns with previous findings that IL-1 induces IL-8 expression and release in keratinocytes and fibroblasts [39, 40]. However, although IL-36γ mRNA and protein expression was induced in HPKs cultivated in SEs upon IL-1 or talabostat treatment (Fig. 4A,C), IL-36γ protein secretion occurred only upon NLRP1 inflammasome activation, which was slightly but significantly reduced upon anakinra co-stimulation (Fig. 5B). Since these results suggest a regulation of IL-36γ secretion by NLRP1 activation, we addressed this hypothesis through 2D monoculture experiments with HPKs. Since IL-1, downstream of NLRP1 activation, induces expression of IL-36γ (Fig. 4B,D), we primed HPKs from 3 different donors with IL-1α to maintain consistently high levels of IL-36γ expression. As both IL-1α and IL-1β act on IL-1R1, we chose to prime with IL-1α to avoid interference with subsequent IL-1β measurements in the supernatant. NLRP1 inflammasome activation by UVB, talabostat, or anisomycin in HPKs revealed secretion of IL-36γ dependent on NLRP1, ASC and GSDMD expression (Fig. 5C,D). This experiment demonstrates that release of IL-36γ, which lacks a signal peptide for canonical secretion, is regulated by NLRP1 activation and occurs most likely through GSDMD pores.

In keratinocytes, cathepsin S is the principal activator of IL-36γ [8]. HPKs constitutively express cathepsin S, although its expression is further increased by IL-1β and particularly by IFN-γ (Fig. 4B,D). This suggests that, in HPKs, IL-36γ is constitutively activated by cathepsin S (Fig. 4D and Fig. 5C,D). In order to characterize cathepsin S expression in the context of NLRP1 activation in HPKs and its relation with IL-36γ, we primed HPKs from 3 different donors with IL-1α (for induction of expression of IL-36γ) and IFN-γ (for induction of expression of cathepsin S), before NLRP1 activation. Interestingly, we detected a release of processed cathepsin S (but not of the full-length protein) dependent on NLRP1, ASC and GSDMD suggesting that active cathepsin S is released together with its substrate IL-36γ through GSDMD pores (Fig. S4A,B).

In conclusion, these results demonstrate that IL-36γ production is regulated by the NLRP1 inflammasome in HPKs. IL-1 secreted upon NLRP1 activation induces expression of IL-36γ and to a lesser extent cathepsin S, before both proteins are released through GSDMD pores.

### NLRP1 activation in SEs induces a molecular signature and histology in the epidermis resembling psoriasis

As expression and secretion of IL-36γ, a cytokine associated with psoriasis and particularly with GPP [7], is strongly induced upon NLRP1 activation in SEs, we compared gene expression of keratinocytes in SEs after NLRP1 inflammasome activation with keratinocytes in lesional psoriasis (Fig. 6A) and in atopic dermatitis as control (Fig. 6B) [41, 42]. Gene set enrichment analysis (GSEA) revealed a common regulation of genes and pathways in the epidermis of talabostat-treated SEs with keratinocytes in psoriasis but not in atopic dermatitis (Fig. 6A,B and Fig. S4C). Most importantly, histological and immuno-histochemical analysis revealed that the epidermal morphology of talabostat-treated SEs shares features known from psoriatic lesions (Fig. 6C). These lesions are characterized by the abnormal retention of nuclei in keratinocytes of the *stratum corneum*, termed parakeratosis. Parakeratosis is a consequence of an increased proliferation and differentiation rate of keratinocytes, leading to a thinning of the stratum granulosum and a thickening of the stratum corneum. Talabostat treatment of SEs induced parakeratosis, which was absent in mock-treated SEs (Fig. 6C). The keratin-binding protein filaggrin is expressed in the granular layer and its expression is dramatically reduced upon talabostat treatment in SEs as well as in lesional psoriasis, further confirming a psoriatic phenotype. Additionally, expression of SPRR2, a marker for keratinocyte differentiation, is strongly induced in psoriatic lesions and upon talabostat treatment in SEs.

**Fig. 6:**
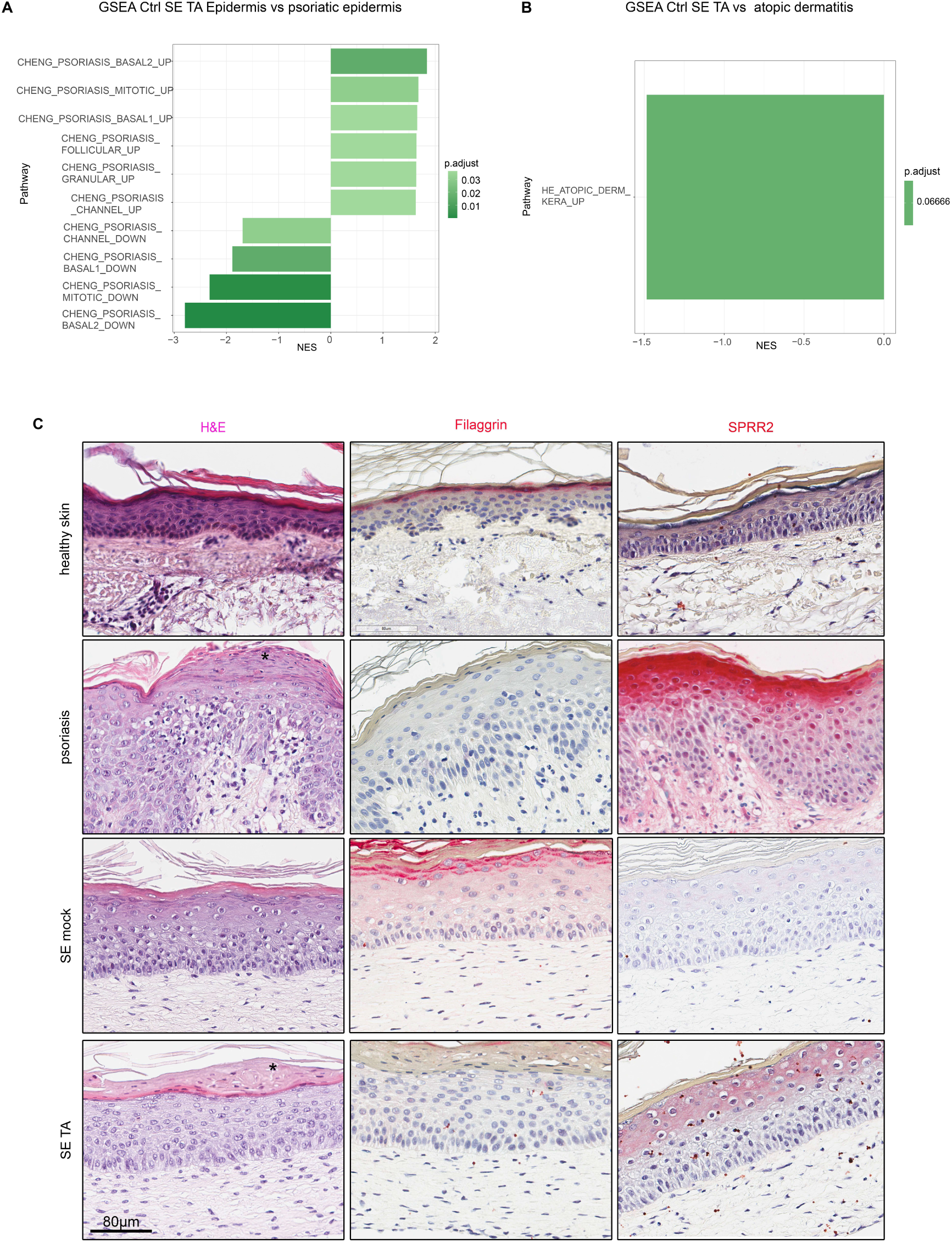
NLRP1 activation in SEs induces a phenotype resembling psoriasis. Talabostat-treated SEs with control HPKs were compared to psoriatic lesions at the mRNA and histological level. GSEA was performed comparing mRNA expression of control HPKs in SEs treated with talabostat (0.3 µM, 3 days) with single-cell RNA sequencing data from **(A)** psoriatic epidermis [41] or **(B)** atopic dermatitis [42]. Gene sets, which were significantly upregulated in psoriatic keratinocytes (positive NES), were also upregulated in the epidermis of control SEs treated with talabostat and vice versa. There was no significant enrichment for atopic dermatitis. **(C)** Histological comparison of healthy and psoriatic skin and mock- and talabostat-treated SEs with control HPKs. H&E, filaggrin and SPRR2 staining. Parakeratosis is indicated (*). Scale bar 80 µm. Stainings are representative of at least 3 independent experiments (SEs) and 3 donors of healthy and psoriatic skin. GSEA: gene set enrichment analysis; TA: talabostat

Although talabostat-treated SEs exhibit several characteristic features of psoriatic skin, they did not develop acanthosis, a typical thickening of the *stratum spinosum* observed in psoriatic lesions. The absence of acanthosis was also reported from other psoriasis models based on SEs and is most likely a consequence of the use of serum-reduced medium, negatively affecting cell proliferation [43].

These data demonstrate that NLRP1 activation in a physiological 3D skin model based on human primary cells induces a molecular signature and a histological phenotype resembling the epidermis in psoriatic lesions in several aspects.

### Inflammasome activation in lesions of patients suffering from psoriasis

It is well accepted that the IL-1 family members IL-1β and particularly IL-36γ play important roles in the pathogenesis of psoriasis [7, 15]. In this study, we demonstrate for the first time that release of IL-36γ, which lacks a signal peptide, is regulated by NLRP1 inflammasome activation in HPKs (Fig. 5D). Furthermore, NLRP1 activation in SEs induced release of IL-17 and IL-23, which are also important for the pathophysiology of psoriasis (Fig. S5). As it is believed that keratinocytes represent the main source of IL-36γ in psoriasis, it is reasonable to hypothesize that inflammasome activation occurs in keratinocytes in active psoriasis. All IL-1-generating human inflammasomes induce the formation of oligomers of the adaptor protein ASC that in turn recruits caspase-1, thereby inducing its activation [16]. Therefore, we determined ASC speck formation in tissue sections of healthy skin and in lesions of patients suffering from psoriasis. In human N/TERT-1 keratinocytes *in vitro*, UVB irradiation induced ASC speck formation in about 1% of the cells, and specks in mock-treated keratinocytes were almost completely absent [23]. Similarly, we identified ASC-positive cells in psoriatic lesions and significantly less in healthy skin (Fig. 7A). Therefore, as suggested by our experiments with SEs, inflammasome activation in active psoriasis occurs in keratinocytes and, most likely, this pathway plays an important role in the pathogenesis of the disease. Although NLRP1 is considered the main inflammasome sensor in human keratinocytes *in vitro* and in human skin [20], NLRP3 or AIM2 expression by keratinocytes or other cell types might also contribute to skin inflammation in psoriasis [15, 44, 45]. To address this point, we analyzed expression of inflammasome sensors, inflammasome proteins, and inflammasome cytokines at the mRNA level in healthy skin and lesional psoriasis. As expected [46], we identified high levels of NLRP1 mRNA but not of NLRP3 or AIM2, and a significant increase in IL-1β, IL-1α, and particularly IL-36γ mRNA in psoriasis compared to healthy skin (Fig. 7B). The expression of NLRP1 in keratinocytes of healthy skin and lesional psoriasis was confirmed by RNAscope, whereas significant levels of NLRP3 and AIM2 could not be detected (Fig. 7C).

**Fig. 7:**
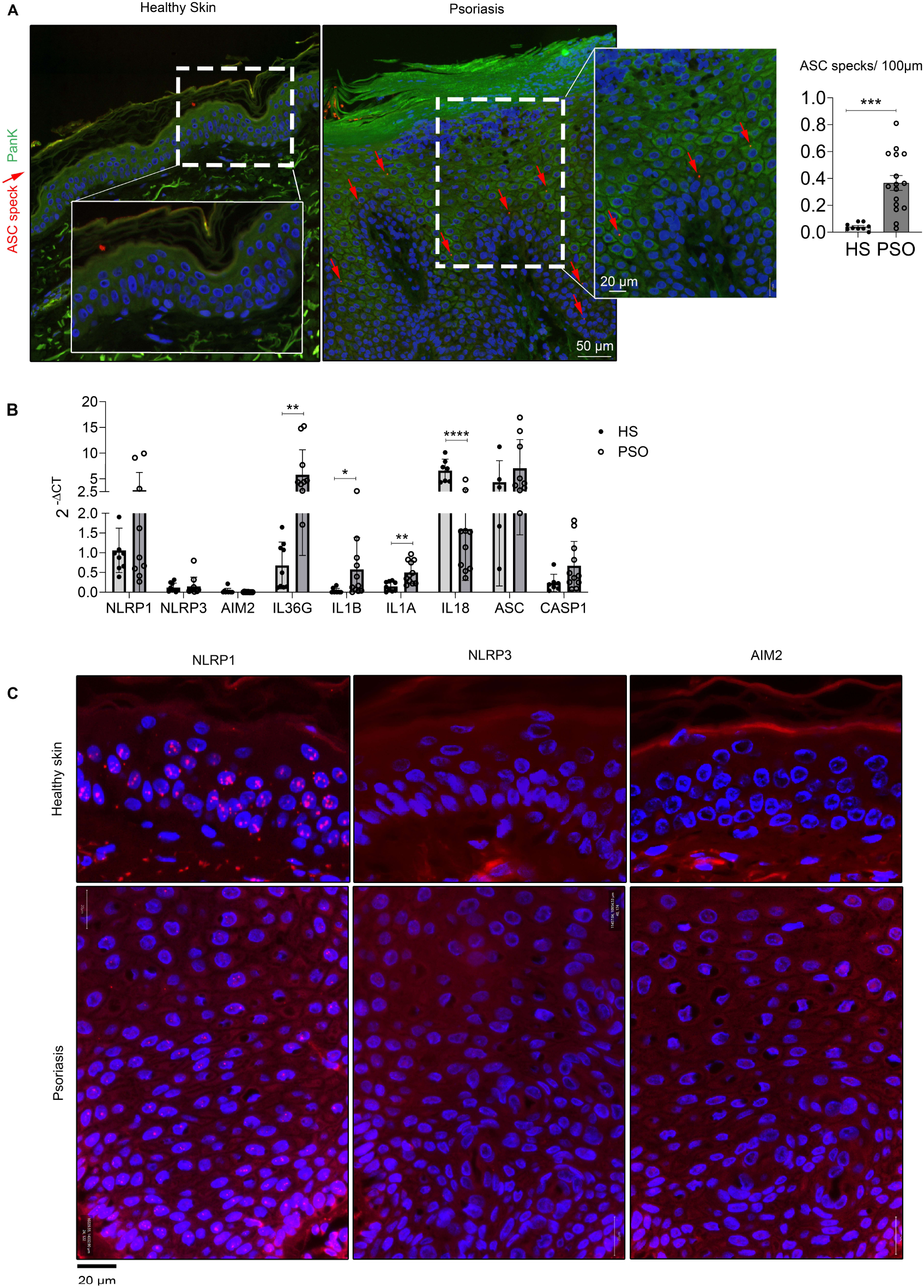
Inflammasome activation and NLRP1 expression in lesional psoriasis. **(A)** Tissue sections of healthy skin or lesional psoriasis were stained for ASC (red), pan-keratin (green) and nuclei (with DAPI, blue). Merge pictures are shown. ASC specks are indicated by arrows and were quantified. Scale bar 50 or 20 µm. Stainings are representative of 9 donors of healthy skin and 16 donors of psoriasis. **(B)** RNA was extracted from FFPE blocks of different donors of healthy skin or lesional psoriasis and expression of the indicated genes was quantified by RT-qPCR. **(C)** RNAscope for NLRP1, NLRP3 and AIM2 (red dots) was performed with biopsies of healthy donors or those suffering from psoriasis. Nuclei are stained with DAPI (blue). Representative merged pictures are shown. Scale bar 20 µm. **(A,B)** P values were calculated with two-tailed unpaired t-test (∗∗∗∗P < 0.0001, ∗∗∗P ≤ 0.001, ∗∗P ≤ 0.01, and ∗P ≤ 0.05, ns = not significant). HS: healthy skin; PSO: psoriatic lesion.

These observations demonstrate that inflammasome activation, most likely through NLRP1, occurs in suprabasal keratinocytes within psoriatic lesions.

### Endogenous dsRNA activates NLRP1 in HPKs

NLRP1 is activated upon viral infection, either through proteolytic cleavage by viral 3C proteases or binding of long viral dsRNA, or through phosphorylation downstream of the RSR, induced by UVB or toxins [22, 26, 27]. However, these stress signals are not relevant for psoriasis. As our results suggest an activation of the NLRP1 inflammasome by keratinocytes in psoriasis, we wondered how NLRP1 might be activated in this inflammatory disease. There is increasing evidence for roles of endogenous dsRNA as a stress factor in several inflammatory conditions [47]. In healthy cells, dsRNA is restricted to the nucleus and mitochondria, but under stress conditions, dsRNA is released to the cytoplasm and extracellular space and considered as new DAMP/HAMP [48]. In the skin, dsRNA is released upon cell damage and contributes through activation of TLR3 to inflammation and repair [49, 50]. Release of the small U1 spliceosomal RNA (U1 RNA), a component of a nuclear ribonucleoprotein complex, has been reported in keratinocytes upon skin injury and is discussed as a stress factor in psoriasis [51]. Furthermore, the amount of total dsRNA is increased in psoriasis due to reduced adenosine-to-inosine (A-to-I) RNA editing [52]. Therefore, we examined RNA localization in healthy skin and lesional psoriasis. Consistent with a previous report [51], we detected cytoplasmic accumulation of RNA in psoriatic keratinocytes, in contrast to its nuclear localization in healthy skin (Fig. 8A). Long dsRNA, such as synthetic poly(I:C), is a known activator of NLRP1 through direct binding of NLRP1 [27]. However, U1 RNA is only 164 base pairs in length and cannot activate NLRP1 upon a physical interaction [27]. Surprisingly, when HPKs were pre-treated (primed) with the TLR3 agonist poly(I:C) before transfection, U1 RNA induced NLRP1 activation, reflected by IL-1β activation and secretion, and the induction of pyroptosis (Fig. 8B). Therefore, U1 RNA, which is discussed as a stress factor in psoriasis, and possibly other short endogenous dsRNAs represent novel activators of NLRP1. These results suggest that NLRP1 is activated in keratinocytes of lesional psoriasis through sensing of endogenous dsRNA such as U1 RNA. The underlying molecular mechanisms are unknown, but it is unlikely that NLRP1 is activated by direct binding of U1 RNA [27].

**Figure 8:**
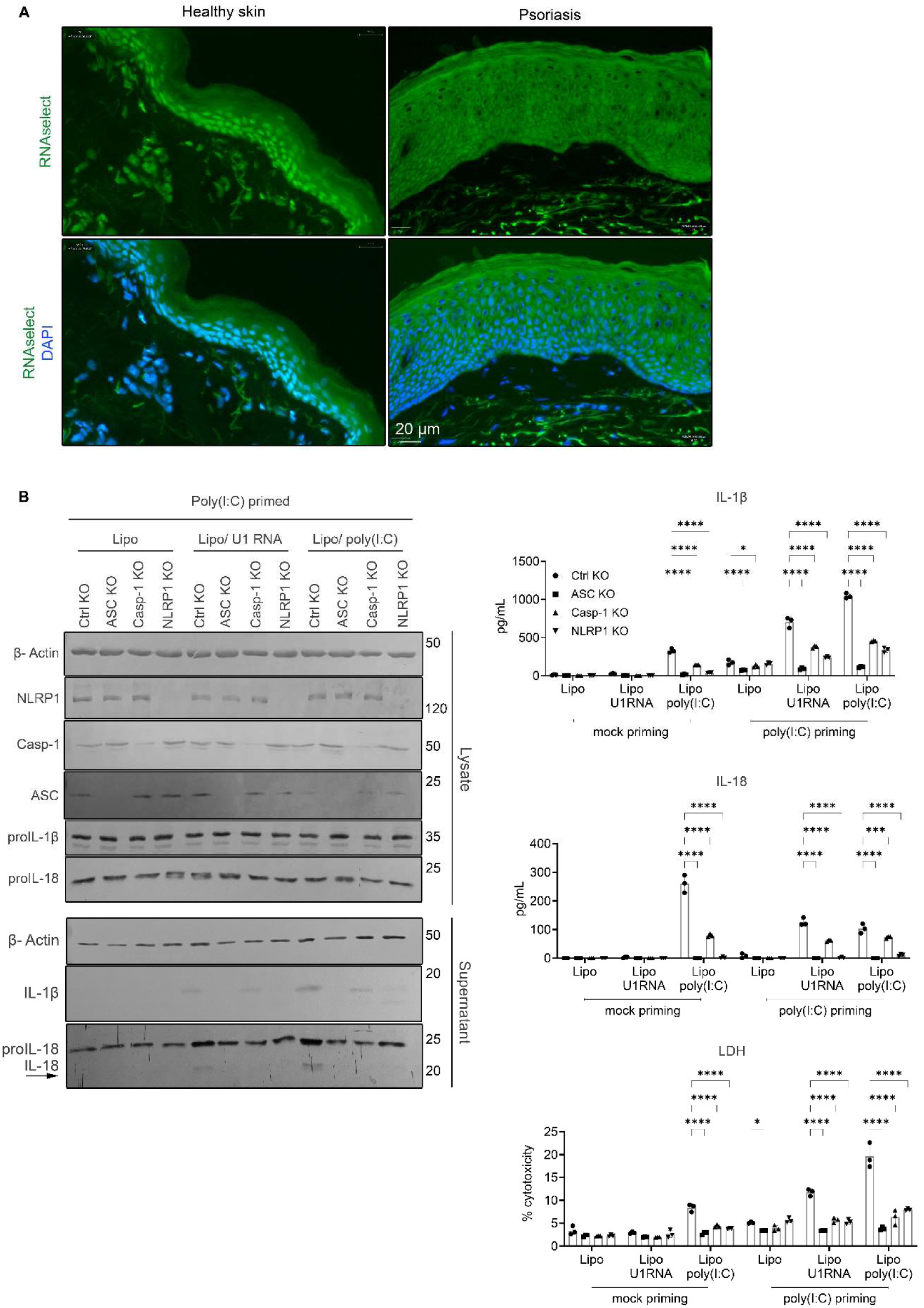
U1 RNA activates the NLRP1 inflammasome. **(A)** Tissue sections of healthy skin or lesional psoriasis were stained for RNA with SYTO RNASelect (green) and nuclei with DAPI (blue). Scale bar 20 µm. Stainings are representative of 3 donors of healthy skin and 3 of psoriasis. **(B)** Western blot for expression of the indicated proteins by polyclonal knockout HPKs from 3 different donors in 2D monolayer. Representative blot out of 3 is shown. HPKs were starved, primed overnight with poly(I:C) (1μg/ml) or mock-treated and then transfected with Lipofectamine (Lipo), Lipo/U1 RNA (1 μg/ml) or Lipo/poly(I:C) (1 μg/ml), for 8 h. Protein expression and release was determined by western blot (left panel) or ELISA (right panel). Cell death was measured by LDH release (right panel). **(B)** Data are represented by mean ± SD of 3 independent donors and are representative of 3 independent experiments. P values were calculated by two-way ANOVA (∗∗∗∗P < 0.0001, ∗∗∗P ≤ 0.001, ∗∗P ≤ 0.01, and ∗P ≤ 0.05, ns = not significant).

## DISCUSSION

Here, we present evidence highlighting a critical role of NLRP1 inflammasome activation in keratinocytes in the pathogenesis of psoriasis. Key findings include the detection of inflammasome activation in lesional psoriasis, accompanied by elevated expression of the inflammasome sensor NLRP1. Using an organotypic skin model, we demonstrate that pharmacological activation of NLRP1 induces histological and molecular changes characteristic for psoriasis. Additionally, supported by 2D monoculture experiments, we revealed a regulation of the IL-36 pathway, which is critically involved in psoriasis, downstream of the NLRP1 inflammasome. Furthermore, we identified NLRP1 activation by endogenous short double-stranded RNA, a factor discussed as a source of stress in psoriasis. We propose that this novel mode of NLRP1 activation is relevant not only for psoriasis but also for other NLRP1-associated inflammatory conditions, offering insight into how NLRP1 functions as a sensor for disturbances of cellular homeostasis, triggering inflammation.

The association of genetic variants, either SNPs or loss-of-function muations, of *IL-1RN* with psoriasis or pustular skin inflammation, demonstrates a causal relationship between IL-1 activity and psoriasis [10, 12, 13]. More specifically, IL-1β is thought to be the key IL-1R1 ligand in psoriasis, for example, by promoting T helper 17 cell maturation [15, 53]. Whereas case reports suggest therapeutic efficacy of IL-1(β) targeting for the treatment of GPP, comprehensive studies are missing [54]. As IL-1β requires activation by caspase-1 downstream of inflammasome activation, this implies critical functions of inflammasomes in psoriasis. Indeed, activation of different types of these protein complexes in different cell types has been described [15, 44, 45]. However, although SNPs of *NLRP1* are associated with psoriasis and NLRP1 is considered the main inflammasome in human skin expressed by keratinocytes, which also produce high levels of IL-1β, a role for NLRP1 and keratinocytes in psoriasis was merely discussed, as exerimental evidence is lacking [20, 21]. In contrast to IL-1β, targeting of IL-36γ is approved for the treatment of patients suffering from GPP [7, 55, 56]. Keratinocytes are the main source of IL-36γ in lesional psoriasis and the main cell type expressing IL-36R [7]. However, to our knowledge, a possible connection between the inflammasome and IL-36γ in keratinocytes has not been explored before. Talabostat treatment of HPKs cultivated in SEs revealed a regulation of IL-36γ production downstream of the NLRP1 inflammasome. Whereas HPKs constitutively express IL-1β, IL-36γ expression must be induced and IL-1β is a very potent inducer of its expression. Most importantly, although IL-1 stimulation is sufficient to induce mRNA and protein expression of IL-36γ in SEs, IL-36γ secretion requires GSDMD prores and, therefore, NLRP1 inflammasome activation. Furthermore, processed cathepsin S, which activates IL-36γ in human keratinocytes in a constitutive manner [8], is also released in a GSDMD-dependent manner, suggesting the intracellular processing of IL-36γ by cathepsin S. These findings demonstrate a regulation of IL-36γ in HPKs by NLRP1 inflammasome activation. In contrast to IL-1β, which is activated and secreted dependent on caspase-1 downstream of NLRP1 in HPKs, IL-36γ is regulated on the transcriptional level by IL-1 and at the level of secretion by GSDMD, both upon NLRP1 inflammasome activation, whereas activation by cathepsin S seems to occur in a more constitutive manner.

Surprisingly, although immune cells are absent in SEs, talabostat-induced activation of NLRP1 triggered a strong pro-inflammatory response, particularly in the dermis by fibroblasts. These cells are often only considered as producers of extracellular matrix molecules, but they have an extraordinary plasticity [57]. Comparison of epidermal gene expression in NLRP1-activated SEs with psoriasis revealed activation of the same pathways. This is supported by histological characterization of SEs, which share characteristic features with psoriasis. Therefore, talabostat-treated SEs represent a novel *in vitro* model of the inflammatory disease based on human primary skin cells. Other psoriasis models based on various types of SEs have been described, where a psoriatic phenotype is induced through different combinations of pro-inflammatory cytokines primarly derived from immune cells associated with psoriasis, the addition of activated T cells, or the culture of skin cells from psoriatic patients [58, 59]. As T cells, cytokines, such as IL-17 or TNF-α, and psoriasis patient-derived skin cells are undoubtedly related to psoriasis, it is obivous to conclude that also NLRP1 activation in keratinocytes is important for inflammation in psoriasis.

The detection of ASC specks in the epidermis of lesional psoriasis represents a direct proof for inflammasome activation. ASC specks are a cardinal sign for activation of all types of inflammasomes, except the CARD8 inflammasome [16]. Although NLRP1 activation in human keratinocytes in 2D culture suggests an activation in basal keratinocytes in the epidermis, talabostat activates NLRP1 in suprabasal cells in SEs and in human skin *ex vivo,* as well as in lesional psoriasis. These results further demonstrate that talabostat-induced NLRP1 activation in SEs represents an excellent model for psoriasis. Other studies suggested an activation of the NLRP3 or AIM2 inflammasome in psoriasis [15, 44, 45]. We detected high levels of NLRP1 mRNA in healthy human skin and lesional psoriasis, but absent or very low expression of NLRP3 and AIM2. This finding suggests, as claimed recently [20], that NLRP1 is the central inflammasome in human keratinocytes, in human skin, and in psoriasis. However, it does not exclude the possibility that activation of other types of inflammasomes in keratinocytes or/and other cell types contributes to inflammation in psoriasis, particularly, when the disease is more systemic [44].

Psoriasis is a very complex genetic disease and it has been discussed for decades what might trigger and initiate cutaneous inflammation [1, 3]. The release of endogenous dsRNA from the nucleus and mitochondria to the cytoplasm and extracellular space can activate several dsRNA-binding receptors, such as RIG-I, MDA5, TLR3, and possibly also NLRP1 [47]. The release of self-RNA induced by cellular stress and sensed by pathways originally known for their anti-viral activities is increasingly suspected to contribute to several inflammatory diseases [60–62], including psoriasis [63, 64]. We confirmed the release of nuclear RNA to the cytoplasms in lesional psoriasis [51]. Furthermore, the total cellular content of dsRNA is increased in psoriasis as a consequence of reduced A-to-I RNA editing [52]. TLR3 activation by dsRNA and the short nuclear U1 RNA play particularly important roles in sunburn, skin injury and skin regeneration [49–51, 65]. Indeed, we could demonstrate NLRP1 activation in HPKs by the short U1 RNA upon priming with the TLR3 agonist poly(I:C). The fact that HPKs sense long dsRNA without priming indicates the activation of another pathway upon TLR3 stimulation required for NLRP1 activation by short dsRNA. NLRP1 binds only long dsRNA suggesting that NLRP1 senses U1 RNA in an indirect manner [27]. Our finding supports the hypothesis of a critical role of NLRP1 in keratinocytes for psoriasis via sensing of endogenous dsRNA, a stress factor, which is associated with the disease [63, 64]. Furthermore, it demonstrates for the first time that NLRP1 can sense cellular perturbations via endogenous short dsRNA.

In summary, our results suggest a critical role of the NLRP1 inflammasome in keratinocytes as a driver and, therefore, as a therapeutic target in psoriasis. Pharmacological inhibition of NLRP1 may provide a dual benefit by simultaneously targeting the IL-1β and IL-36γ pathways. NLRP1-activated SEs represent a novel model for psoriasis, which might contribute to the development of NLRP1 inhbitors. NLRP1 activation takes place in suprabasal keratinocytes of the psoriatic epidermis, which is highly accessible for topical treatment. This strategy might not only control local inflammation effectively, but also improve the overall safety profile of psoriatic treatments by limiting drug activity to the skin.

## MATERIALS AND METHODS

### Human skin samples

Surplus fresh or paraffin-embedded skin biopsies were collected from plastic-surgery patients and psoriatic patients (Dermatology, University Hospital Zürich) with informed written consent, following approval from the Kantonal Ethics Committee (KEK) of Zürich (KEK-ZH-Nr. 2015-0198 and BASEC-Nr. 2024-01030). All procedures were conducted in accordance with the Declaration of Helsinki principles.

### Culture and treatment of primary cells in 2D

Isolation and culture of human primary keratinocytes (HPKs) was performed as described previously [27, 71]. HPKs were grown in keratinocyte serum-free medium (K-SFM, Thermo Fisher Scientific, USA) supplemented with epidermal growth factor (EGF) and bovine pituitary extract (BPE), harvested in trypsin/EDTA solution (0.05%/0.02% w/v) and either cultured for at least 48 hours before experiments, or used to generate the epidermis of fibroblasts-derived matrix-based skin equivalents.

Human dermal fibroblasts (HDFs) were isolated from dermis upon incubation with a solution of 1 mg/ml collagenase (Sigma) and 0.05 mM CaCl2 in phosphate buffered saline for 2 hours at 37°C. A single cell suspension was obtained by pipetting the dermis up and down in DMEM (high glucose, pyruvate, Thermo Fisher Scientific) containing 25% fetal bovine serum (PAN-Biotech, Germany) and 1% antibiotic/antimycotic (Thermo Fisher Scientific).

HDFs were grown in DMEM supplemented with 10% FBS (Sigma) and 1% antibiotic/antimycotic, harvested in trypsin/EDTA solution (0.05%/0.02% w/v) and either cultured for at least 48 hours before experiments, or used to generate the fibroblasts-derived matrix-based dermal equivalents.

HPKs or HDFs in monolayer were stimulated in KSFM pure (Thermo Fisher Scientific) with talabostat (0.3 µM or 10 µM, Lucerna-chem, HY-13233A), recombinant human (rh) IL-1β (10 ng/ml, PeproTech 200-01B), rh IL-18 (20 ng/ml, Gibco PHC0186), rh IL-36γ (100 ng/ml, R&D 6835-IL), rh TNF-α (10 ng/ml, Invivogen rcyc-htnfa), rh IFN-γ (20 ng/ml, Invivogen rcyec-hifng), poly(I:C) (1 µg/ml, InvivoGen tlrl-pic), LL-37 (1 µg/ml, InvivoGen tlrl-l37), anisomycin (1 µM, Sigma A9789) and UVB (86.4 mJ/cm^2^, UV802L, WALDMANN, Villingen-Schwenningen, Germany). K-SFM containing the corresponding amount of DMSO or PBS served as control.

### Generation of CRISPR/Cas9 knockout keratinocytes by Lentivirus transduction

CRISPR/Cas9-mediated gene knockout was performed as described [71]. Briefly, single-stranded DNA oligonucleotides were designed on the Benchling platform (https://benchling.com) and cloned into pLenti CRISPR v2 plasmid (sgRNA sequences in Expanded View Table 1). The pLenti CRISPR v2 containing the specific sgRNA was co-transfected into HEK 293T cells with the envelope and packaging plasmids psPAX2 (#12260, Addgene) and pMD2.G (#12259, Addgene). The supernatant containing viral particles was harvested and filtered (Sartorius Minisart 0.45 μm, Thermo Fisher Scientific) 48 h later.

HPKs were transduced after being isolated from fresh biopsiesm, while co-cultured on a layer of antibiotic-resistant feeder cells. Selection of keratinocytes was performed with puromycin 5 μg/ml in the presence of antibiotic-resistant feeder cells [71]. Knockout efficiency was assessed at the protein level by western blot. Lentivirus-transduced knockout keratinocytes were used for the generation of skin equivalents.

### Generation of CRISPR/Cas9 knockout keratinocytes by electroporation

CRISPR/Cas9-mediated gene knockout was performed as described [72]. Briefly, RNP complexes were prepared by mixing Cas9 protein with custom-designed sgRNA (https://benchling.com) (in a 1:1 molar ratio https://chopchop.cbu.uib.no) (sgRNA sequences in Expanded View Table 1), followed by incubation at room temperature for 20 minutes. Cell pellets were resuspended in electroporation buffer R (5 μl buffer R per 0.15 × 106 cells) and then transferred to the RNP mix. Electroporation was performed according to the manufacturer’s protocol, carried out at 1700 V/20 ms/1 pulse. Cells were seeded into flasks and cultured for 1 week post-electroporation before further experiments.

### Generation of fibroblast-derived matrix-based dermal equivalents

Dermal equivalents were generated as described [35]. Briefly, HDFs of passage 5 were expanded for a maximum of three passages in DMEM supplemented with 10% FBS (Sigma) and 1% antibiotic/antimycotic. 5 x 105 HDFs were seeded on day 1, 3 and 5 (in total 1.5 x 106) onto 12-well translucent ThinCerts with high-density 0.4 μm pores (11.31 cm2 culture area, VWR, Switzerland), placed in deep-well plates (12-well plate, VWR) and incubated at 37°C, 5% CO2 and 20% O2 for another 3 weeks. The final input of HDFs was 1.5 x 106 per transwell insert. HDFs were cultivated in 3:1 DMEM/Ham’s F12 (FAD) containing 10% FBS (Sigma) and 1% antibiotics/antimycotics, supplemented with 2-phospho-l-ascorbic acid (200 μg/ml - Sigma) and the additional recombinant human proteins: TGFβ-1 (1 ng/ml - Thermo Fisher Scientific), EGF (2.5 ng/ml - Thermo Fisher Scientific), b FGF (5 ng/ml-Peprotech), and insulin (5 μg/ml - Sigma). The medium was changed every two days for the duration of the 4-week period.

### Generation of fibroblasts-derived matrix-based skin equivalents

Skin equivalents (SEs) were generated as described [35]. Briefly, 2.5 x 105 HPKs, wild type or CRISPR/Cas9-modified, were seeded per dermal equivalent in FAD medium containing 10% FBS and 1% antibiotics/antimycotics, supplemented with 2-phospho-l-ascorbic acid-trisodium salt (200 μg/ml), hydrocortisone (0.4 μg/ml - Sigma), and cholera toxin (1×10−10 M - Sigma). After 3 days of submerged growth, the skin equivalents were air-lifted and the medium changed every other day, for 2 weeks.

### Treatment of skin equivalents

Skin equivalents, 10 days from the airlift, were stimulated with Opti-MEM containing talabostat (0.3 or 10 μM), anakinra (10 μg/ml - Swedish orphan biovitrum, Sweden) or rh IL-1β or rh IL-1α (1 ng/ml - Peprotech).

### Transfection of keratinocytes in monolayer

Keratinocytes were transfected with Lipofectamine 2000 Transfection Reagent (ThermoFisher – 11668027) according to manufacturer’s instruction. Briefly, 1 µg of U1 RNA or poly(I:C) was diluted in 100 µl of K-SFM pure, per well, and mixed gently. Separately, 2 µl of Lipofectamine 2000 was diluted in 100 µl of K-SFM pure, per well. The diluted Lipofectamine 2000 was incubated at room temperature for 5 minutes. After the incubation, the diluted RNA was combined with the Lipofectamine 2000 solution (total volume 200 µl per well). The RNA-Lipofectamine solution was gently mixed by pipetting and incubated at room temperature for 15 minutes. The transfection mixture was added dropwise to each well, containing 800 µl of fresh K-SFM medium and incubated at 37°C in a 5% CO₂ incubator for 8 hours.

### Real-time PCR

Skin equivalents were homogenized using the TissueLyser II (3 minutes, 30/s Hz, QIAGEN). Total RNA from monolayer keratinocytes or homogenized skin equivalents was isolated using Trizol (Qiagen, Germany), according to the manufacturers’ instructions. Levels of mRNA were determined by quantitative real-time PCR using the LightCycler 480 instrument and FastStart Essential DNA Green Master (Roche, Switzerland) and specific primers (Microsynth, Switzerland). mRNA levels were normalized to HPRT and RPL27. Primers can be found in Expanded View Table 2.

### Immunoblotting

Monolayer cell lysates were harvested with SDS loading buffer and protein amounts were estimated by β-actin band quantification after immunoblotting. Epidermis and dermis of the skin equivalents were collected separately upon dispase II digestion (30 minutes, 37 °C) and lysed in a buffer containing 4% SDS and 100 mM Tris HCl pH 7.6, with TissueLyser II (3 min, 30/s Hz, QIAGEN). Proteins in solution were then incubated at 95°C for 5 minutes, sonicated and separated from the pellet upon centrifugation (16.000 g, 5 minutes). Protein concentration was measured with the BCA protein assay (PIERCE, USA). Before loading in the gel, 0.1 M DTT was added to the mix and proteins were diluted 1:5 in 5X loading buffer (100% glycerol + 1% bromophenol blue).

Cell culture supernatants were precipitated with 2.5 volumes of acetone (100% w/v, Sigma-Aldrich, USA) by overnight incubation at −20°C followed by centrifugation for ca. 2 h (4,000g at 4°C) and resuspended in SDS loading buffer. Proteins were separated by SDS PAGE and analyzed by immunoblotting as previously described [27]. The primary and secondary antibodies used are specified in Expanded View Table 3.

### Histology

Human tissue or skin equivalents samples were fixed overnight in formalin 4%, dehydrated automatically and embedded in paraffin (Leica EG 1150, Leica microsystem). Paraffin blocks were cut into 5 μm-thick sections and stained with hematoxylin and eosin (H&E) according to standard protocols or analyzed by immunohistochemistry or immunofluorescence.

### Immunohistochemistry

Specific stainings on paraffin sections were performed using standard immunohistochemistry protocols. Briefly, tissue sections were deparaffinized in xylene and rehydrated by 2 min incubation in decreasing concentrations of ethanol (100% - 96% - 80%-70% and 50%), and 2 min in water. The activity of endogenous peroxidase was blocked by 10 minutes of incubation in 3% peroxidase-blocking solution (hydrogen peroxide 30% in ddH2O; Merck, New Jersey). For antigen retrieval, slides were heated in a steamer for 30 minutes in antigen retrieval solution, pH 9 (Dako, Denmark) or sodium citrate buffer, pH 6.0 (10 mmol/l sodium citrate, 0.05% Tween 20). Sections were then blocked for 1 h at room temperature with 5% bovine serum albumin in PBST (0.05% Tween 20 in PBS, blocking solution) and incubated overnight at 4°C with the primary antibody diluted in the blocking solution. Biotinylated secondary antibodies were applied for 1 h at room temperature. Stainings were visualized with the Vectastain Elite ABC HRP Kit (Vector Laboratories, United States) and the AEC substrate (Dako); nuclei were stained using Mayer’s hematoxylin (Kantonsapotheke, Zurich, Switzerland). Slides were mounted with Faramount aqueous mounting medium (Dako) and scanned using an Aperio ScanScope (Leica Biosystems, Germany) or with the PhenoImager Vectra Polaris (Akoya Biosciences, United States). The primary and secondary antibodies used are specified in Expanded View Table 3.

### Immunofluorescence

Paraffin slides were processed until antigen damasking as described for immunohistochemistry. Then, sections were cooled down, washed in 1x PBST and incubated for one hour in blocking solution. Slides were incubated with the primary antibody diluted in blocking solution overnight at 4 ◦C. The day after, slides were washed three times in PBST and incubated with the Alexa Fluor conjugated secondary antibody (Thermo Fisher Scientific) 1:200 in blocking solution for 1 h at room temperature. Then, the secondary antibody was removed and DAPI (Sigma, 1:1000 in blocking solution) applied to the slides, for 5 minutes at room temperature. After 3 washes in 1x PBST, coverslips were mounted on glass slides using prolong Gold antifade mounting medium (Thermo Fisher Scientific). Tissue slides were visualized using fluorescence microscopy (Zeiss Axiocam 503 mono) and scanned with the PhenoImager Vectra Polaris (Akoya Bioscience, United States). Details of the primary and secondary antibodies used can be found in Expanded View Table 3.

### ELISA and cytotoxicity assay

Cell culture supernatants were collected, cleared by centrifugation (400 g, 3 minutes) and release of human IL-1β, IL-1α, IL-18, IL-36γ and IL-8 was measured by ELISA (R&D Systems or Biolegend (IL-8), USA) according to manufacturer’s instruction. Cytotoxicity was evaluated by measuring the release of lactate dehydrogenase (LDH) in the supernatant using the CytoTox 96 non-radioactive cytotoxicity assay (Promega, Fitchburg, WI). To obtain total LDH, cells were lysed for 30 min in 10% Triton X-100 (Sigma). Results are shown as percentage release (LDH in SN/total LDH).

### Proteome profiler human XL cytokine array

Cytokine array was performed using the supernatant of skin equivalents either mock-, or talabostat-(0.3 μM) or anakinra (10 μg/ml) plus talabostat-treated (3 days), using the proteome profiler human XL cytokine array kit (R&D), according to the manufacturers’ instructions.

### Quantification of ASC specks

Sections of SE, healthy or psoriatic skin were stained for ASC speck formation and entirely scanned with the PhenoImager Vectra Polaris. Sections were analyzed using the Open Software for Bioimage Analysis QuPath: ASC specks were manually counted and the length of the skin section was measured. The number of specks was normalized to a length of 100 μm.

### Quantification of epidermal thickness in SEs

Epidermal thickness was measured over the entire length of section using the software Aperio Image Scope (Leica Biosystem) on H&E scanned slides. Measurements were taken for the living epidermis, from the basal layer to the granulosum.

### High-throughput cDNA sequencing

Three replicates of Ctrl SEs and ASC KO SEs were treated with 0.3 μM talabostat for 3 days. Mock treatment served as control. Epidermis and dermis were separated with dispase II digestion and RNA isolated with RNeasy Plus Universal Mini Kit (QIAGEN), upon homogenization with the TissueLyser II (QIAGEN).

High-throughput sequencing of the coding transcriptome for gene expression profile was performed with the Illumina Novaseq 6000 (Functional Genomics Center Zurich, FGCZ). Library preparation was achieved with approximately 0.1 – 1 μg total RNA, using the Illumina Truseq Stranded mRNA kit, according to the manufacturers’ instructions (FGCZ). The Agilent TapeStation was used for cDNA quantification and quality assessment. 25M reads were analyzed per sample, using a single read 100 bp configuration.

### RNA-Seq Read Mapping

Gene-level counts were quantified from single-end reads aligned to the GRCh38.d1.vd1 genome using STAR v2.7.5a [73].

### Identification of Differentially Expressed Genes (DEGs)

Differential gene expression was performed with limma voom [74] and visualized with ComplexHeatmap [75]. For epidermis analysis, contrasts included ctrl untreated versus ctrl talabostat-treated, Ctrl untreated versus ASC KO untreated, ctrl talabostat-treated vs ASC KO talabostat-treated, ASC KO untreated versus ASC KO talabostat-treated. Same contrasts were analyzed for dermis. Multiple testing correction was performed with the Benjamini-Hochberg procedure. The gene expression data generated in this study were deposited in the Gene Expression Omnibus (GEO) under the accession number GSE282206.

### Functional Analysis of DEGs

Pathway analysis was performed using clusterProfiler (Yu 2012 OMICS: A journal of Integrative Biology) and GSEA [76] with MSigDb and additional pathways from the following papers [46], [47] and [38].

### RNAscope In Situ Hybridization

To detect target RNA expression, we performed RNAscope in situ hybridization using the RNAscope Multiplex Fluorescent Reagent Kit v2 (Advanced Cell Diagnostics, Newark, CA) following the manufacturer’s protocol. Briefly, subsequent tissue sections were pre-treated with the appropriate reagents for protease digestion. Specific RNA probes designed for human-NLRP1, human-AIM2 or human-NLRP3 were hybridized to the subsequent tissue samples, and signal amplification was achieved through a series of hybridization steps involving multiple amplification probes. Fluorescent detection was performed using Opal 520 (Akoya Bioscience). Sections were counterstained with DAPI to visualize cell nuclei. The stained sections were imaged and scanned using the PhenoImager Vectra Polaris (Akoya Bioscience).

### RNA Staining with SYTO® RNASelect™ Green Fluorescent Cell Stain

To visualize RNA within cells, we utilized SYTO® RNASelect™ Green Fluorescent Cell Stain (Thermo Fisher Scientific) following the manufacturer’s instructions. OCT-embedded human skin and psoriatic biopsy samples were sectioned at a thickness of 10 µm using a cryostat. The tissue sections were immediately fixed in pre-chilled methanol at −20°C for 10 minutes to preserve cellular structures, followed by three washes in phosphate-buffered saline (PBS). After fixation, the sections were incubated with 500 nM SYTO® RNASelect™ Green Fluorescent Cell Stain in PBS for 20 minutes at 37°C in the dark to selectively label intracellular RNA. Cell nuclei were counterstained with DAPI. Excess dye was removed by washing the sections thoroughly with PBS. Fluorescent images were captured using a fluorescence microscope equipped with a FITC filter to visualize RNA.

### Statistical analysis

Statistical analysis for not high throughput data was performed using the software Prism (GraphPad Software, La Jolla). For the specific tests, please see figures legends. Differences were considered significant when P values were below 0.05 (∗∗∗∗P < 0.001, ∗∗∗P ≤ 0.001, ∗∗P ≤ 0.01, and ∗P ≤ 0.05, ns = not significant).

## Acknowledgments

We are grateful for generous financial support by the LEO Foundation (LF-OC-23-001156), Wilhelm Sander Foundation (2019.075), Unversity of Zurich, Bruno Bloch Foundation, Huggenberger-Bischoff Foundation, and Swiss National Science Foundation (310030_197426). HDB, MDF, JTM, MPL and ToK are members of the SKINTEGRITY.CH collaborative research consortium. MS and TK are members of the PhD program in Molecular Life Sciences Zürich. We thank Prof. Sabine Werner (ETH Zürich) for continuous support. We are thankful for the support by Federica Sella, Elizabeth Pavez Lorie and Conrad Wyss.

## Availability of Data and Materials

The gene expression data generated in this study were deposited in the Gene Expression Omnibus (GEO) under the accession number GSE282206.

## Author contributions

Conceptualization: MDF, HDB

Methodology: MDF, TK, PFC, PH, HDB

Investigation: MDF, TK, PFC, PH, HDB

Visualization: MDF, PH

Funding acquisition: HDB

Project administration: HDB

Supervision: HDB

Writing – original draft: HDB, MDF

Writing – review & editing: MDF, TK, PFC, PH, MS, PB, SP, IK, MPL, HDB, ToK, JTM

