## Supplementary materials for "NLRP1 inflammasome activation in skin equivalents reveals mechanistic insights into the roles of keratinocytes in psoriasis"

**Supplementary Material**

**TABLES**

**Table S1:**

**sgRNA for lentiviral transduction**

| Non-targeting control (Ctrl.) | |
|---|---|
| Forward oligo | CACCGGTAGCGAACGTGTCCGGCGT |
| Reverse oligo | AAACACGCCGGACACGTTCGCTACG |

| ASC | |
|---|---|
| Forward oligo | CACCGTAGAAGCTGACCAGCTTGT |
| Reverse oligo | AAACACAAGCTGGTCAGCTTCTAC |

| Caspase-1 | |
|---|---|
| Forward oligo | CACCGATTGACTCCGTTATTCCGAA |
| Reverse oligo | AAACTTCGGAATAACGGAGTCAATC |

| NLRP1 | |
|---|---|
| Forward oligo | CACCGCTCAGCCAGAGAAGACGAG |
| Reverse oligo | AAACCTCGTCTTCTCTGGCTGAGC |

**sgRNA for electroporation**

Non-targeting control (Ctrl.): AAATGTGAGATCAGAGTAAT

NLRP1: CTGGATCCATGAATTGCCGG

ASC: CGCTAACGTGCTGCGCGACA

Caspase-1: TCCACTAGCATCTTACCTTG

**Table S2: Real-time PCR primers**

| | |
|---|---|
| CCL3 | |
| Forward | 5'-AGTTCTCTGCATCACTTG CTG-3' |
| Reverse | 5'-CGGCTTCGCTTGGTTAGGAA-3' |
| CCL5 | |
| Forward | 5'-AAGTTGTCTGTGTGCGCAAATCC-3' |
| Reverse | 5'-CCATTCCAGAAAAGCCACAGTTTT-3' |
| CCL20 | |
| Forward | 5'-CCAGCAGTCGTCTTT GTCAC-3' |
| Reverse | 5'-CTCTGGGTTGGCACACACTT-3' |
| DEFB4A | |
| Forward | 5'-GGTGGTATAGGCGATCCTGTT-3' |
| Reverse | 5'-AGGGCAAAAGACTGGATGACA-3' |
| HPRT | |
| Forward | 5'-ATTGTAATGACCAGTCAACAGGG-3' |
| Reverse | 5'-GCATTGTTTTGCCAGTGTCAA-3' |
| IL-1B | |

| | |
|---|---|
| Forward | 5’-CACGATGCACCTGTACGATCA-3’ |
| Reverse | 5’-GTTGCTCCATATCCTGTCCCT-3’ |

| CXCL8 | |
|---|---|
| Forward | 5’-TTTTGCCAAGGAGTGCTAAAGA-3’ |
| Reverse | 5’-AACCCTCTGCACCCAGTTTTC-3’ |

| IL-36γ | |
|---|---|
| Forward | 5`-AGGAAGGGCCGTCTATCAATC-3' |
| Reverse | 5`-CACTGTCACTTCGTGGAACTG-3' |

| MEFV | |
|---|---|
| Forward | 5`-TAAGACCCCTAGTGACCATCT G-3' |
| Reverse | 5`-TTCCCCATAGTAGGTGACCAG-3' |

| LCE3E | |
|---|---|
| Forward | 5’-AGTACAGTGTCTGCCTCCAGCT-3’ |
| Reverse | 5’-CTGTCACAGGAGTTGGACCTCT-3’ |

| LCN2 | |
|---|---|
| Forward | 5’-CCACCTCAGACCTGATCCCA-3’ |
| Reverse | 5’-CCCCTGGAATTGGTTGTCCTG-3’ |

| SPRR2B | |
|---|---|
| Forward | 5`-ACGCCAAAGTGCCCAGAG-3` |
| Rev<br>erse | 5`-ATTTCTGCTGGCACTGCTGAG-3` |

| CTSS | |
|---|---|
| Forward | 5`-GGATCACCACTGGCATCTCT-3` |
| Reverse | 5`-ATTCCCATTGAATGCTCCAG-3` |

| AIM2 | |
|---|---|
| Forward | 5`-CAGAAATGATGTCGCAAAGCA-3' |
| Reverse | 5`-TCAGTACCATAACTGGCAAACAG-3' |

| NLRP3 | |
|---|---|
| Forward | 5`-GCAAAAAGAGATGAGCCGAAGT-3' |
| Reverse | 5`-GCTGTCTTCCTGGCATATCACA-3' |

| CASP-1 | |
|---|---|
| Forward | 5`-TCCCTAGAAGAAGCTCAAAGGATATG-3' |
| Reverse | 5`-CGTGTGCGGCTTGACTTG-3' |

**Table S3: antibodies**

| **Antibody** | **Order number** | **Application** |
|---|---|---|
| β-actin | A5441 (Sigma) | WB |
| ASC | AL177 (Adipogen) | WB, IHC, IF |
| Caspase-1 | sc-622 (Santa Cruz) | WB |
| Filaggrin | NBP1-87528 | IHC |
| IL-18 | PM014 (MBL) | WB |
| IL-1β | MAB201 (R&D) | WB |
| IL-36γ | AP2320 (R&D) | WB |
| CTSS | PA5-47088 (Thermofisher) | WB |

| | | |
|---|---|---|
| GSDMDC1 | NBP2-33422 (Novus) | WB |
| Isotype control mouse | ab37355 (abcam) | IHC, IF |
| Isotype control rabbit | ab172730 (abcam) | IHC, IF |
| Keratin-10 | 905404 (Biolegened) | IF |
| Keratin-15 | GP-CK15 | IF |
| Pan-cytokeratin | sc-8018 | IF |
| SPRR2 | AG-25B-0002 | IHC |
| Anti-Rabbit IgG (Fc), AP Conjugate | s373b (Promega) | Secondary AB, WB |
| Anti-Mouse IgG (H+L), AP Conjugate | s372b (Promega) | Secondary AB, WB |
| Anti-Goat IgG, AP Conjugate | V115A (Promega) | Secondary AB, WB |
| Anti- Guinea pig IgG (H+L), AP Conjugate | ab6909 (Abcam) | Secondar AB, WB |
| Anti-mouse IgG (H+L) Alexa Fluor 488 | A21429 (Life technologies) | Secondar AB, IF |
| Anti-mouse IgG (H+L) Alexa Fluor 647 | A11001 (Thermo Fisher Scientific) | Secondar AB, IF |
| Anti-mouse IgG BIOT | Ab5886 | Secondar AB, IHC |
| Anti-Rabbit IgG BIOT | 4010-08 (Southern Biotech) | Secondar AB, IHC |

### Supplementary Figures

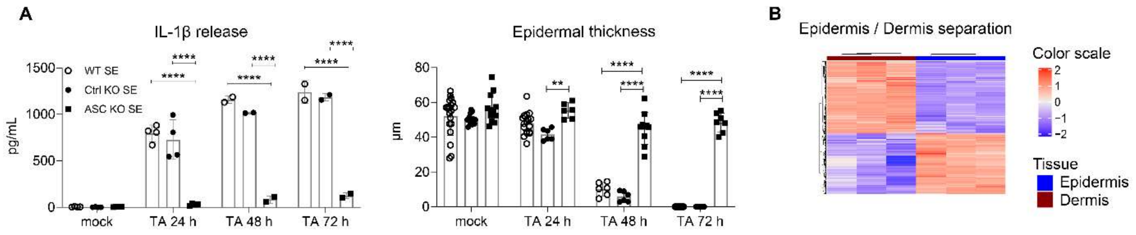


C

| Pathways Epidermis | Leading edge genes |
| --- | --- |
| REACTOME_INNATE IMMUNE_SYSTEM | MUC4, PIGR, MUC16, DEFB4A, DEFB4B, ITGAX, SAA1, SERPINA1, LTF, ALOX5, MUC21, ITGB2, ATP6V0A4, CEACAM6, LCN2, MEFV, NCF2, RNASE7, CFB, IQGAP2, NFAM1, VNN1, NCF4, S100A12, CD177, CEACAM1, MUC20, RAC2, MUCL1, IL1B, ARG1, S100A7A, CTSS,LAT2, CD68, ICAM2,ENPP4, CRISPLD2, PI3 |
| GOBP_DEFENSE_RESPONSE TO_OTHER_ORGANISM | SH2D1B, IL36A, DEFB4A, ITGAX, LCE3C, LTF, OASL, LCN2, MEFV, WFDC2, NCF2, RNASE7, IL23A, LCE3A, CFB, NRROS, VNN1, IL36G, A2M, GPER1, S100A12, CD177, KYNU, CEACAM1, PLA2G2F, SLA, IL1B, SERPINB4, ARG1, GSDMA, CX3CL1, IL36RN, NLRP10, GBP5, SERPINB9, IL36B, PI3, CXCL6, SRMS |
| GOBP_IMMUNE_ RESPONSE | SH2D1B, PIGR, IL36A, DEFB4A, IL1RL1, CSF3, VTCN1, LTF, CXCR4, ALOX5, ITGB2, OASL, LCN2, MEFV, WFDC2, IL19, NCF2, RNASE7, IL23A, CFB, NRROS, NFAM1, VNN1, IL36G, A2M, CSF2RB, GPER1, S100A12, PKN1, CD177, BMP6, KYNU, CEACAM1, KCNN4, SPNS2, RAC2, PLA2G2F, SLA, RAPGEF3, IL1B, SERPINB4, ARG1, CTSS, CX3CL1, IL36RN, LAT2, TNFSF14, CD68, NLRP10, GBP5, SERPINB9, SLAMF9, IL36B, PI3, SMAD6, CXCL6, SRMS, GPRC5B |
| GOBP_BIOLOGICAL_ PROCESS_INVOLVED IN_INTERSPECIES_ INTERACTION BETWEEN_ORGANISMS | TMPRSS2, SH2D1B, REN, IL36A, DEFB4A, ITGAX, PLAAT3, CSF3, VTCN1, LCE3C, LTF, CXCR4, OASL, LCN2, MEFV, WFDC2, NCF2, RNASE7, IL23A, LCE3A, CFB NRROS, TH, VNN1, IL36G, A2M, CSF2RB, GPER1, S100A12, CD177, BMP6, KYNU, CEACAM1,PLA2G2F, SLA, IL1B, SERPINB4, ARG1, DUOX2, GSDMA,TIMP4, CX3CL1, AKAP12, IL36RN, CD68, NLRP10, GBP5, SERPINB9, IL36B, PI3, SMAD6, CXCL6, SERPINB3, SRMS |
| GOBP_DEFENSE_ RESPONSE | SH2D1B, DUOXA2, IL36A, DEFB4A, ITGAX, SAA1, IL1RL1, SERPINA1, LCE3C, LTF, CXCR4, ALOX5, ITGB2, SAA2, OASL, LCN2, MEFV, WFDC2, NCF2, RNASE7, IL23A, LCE3A, CFB ,NRROS, NFAM1, VNN1, IL36G, A2M, GPER1, S100A12, CD177, BMP6, KYNU, LYPD1, CEACAM1, CHST2, KCNN4, GAL, PLA2G2F, SLA, CAMK2N1, IL1B, SERPINB4, ARG1, DUOX2, GSDMA, CX3CL1, IL36RN, MMP3, CD68, NLRP10, GBP5, SERPINB9 |
| GOBP_INNATE_ IMMUNE_RESPONSE | SH2D1B, IL36A, LTF, OASL, LCN2, MEFV, WFDC2, NCF2, RNASE7, IL23A, CFB, NRROS, VNN1, IL36G, A2M, GPER1, S100A12, CD177, KYNU, CEACAM1, PLA2G2F, SLA, SERPINB4, ARG1, CX3CL1, IL36RN, NLRP10, GBP5, SERPINB9, IL36B, PI3, SRMS |
| GOBP_EPITHELIAL_ CELL_DIFFERENTIATION | SPRR2F, ROBO4, PLAAT3, LCE3C, CXCR4, PLAAT4, SLC44A4, LCE3E, XDH, SPRR3, KRT13, SPRR2A, DHRS9, LCE3D, STC1, LCE3A, KRT19, SPRR2B, FOXA1, SPRR2E, AGR2, RHCG, SPRR2D, BMP6, SPRR2G, CEACAM1, RAPGEF3, IL1B, CNFN, TJP3, DAB2, HEY1, TGM1, KRT23, ID3, ALOX15B, ATOH8, CDH2, TIE1, DLX3, LAMA1, PDE2A, SULT1B1, CDSN, CES1, LCE1F, HPSE, POF1B |
| GOBP_INFLAMMATORY RESPONSE | DUOXA2, IL36A, SAA1, IL1RL1, SERPINA1, CXCR4, ALOX5, ITGB2, SAA2, MEFV, IL23A, NRROS, NFAM1, VNN1, IL36G, A2M, GPER1, S100A12, BMP6, CHST2, GAL, CAMK2N1, IL1B, CX3CL1, IL36RN, MMP3, CD68, NLRP10, GBP5, IL36B, CXCL6, GPRC5B, C3, CXCL5,GBA, ADGRE5 |
| GOBP_CENTRAL_NERVOUS_ SYSTEM_DEVELOPMENT | MAPT, GJC2, RECK, SOX6, SLIT3, HOXB3, CTNNA2, WNT4, S1PR1, FOXP2, ROBO2, NHLH2, LOXL3, ASPA, PCSK1, GLI2, SCN1B, SELENOP, PCDH18, PLXDC1, SLC8A1, SHROOM4, CLU |

| Pathways Dermis | Leading edge genes |
| --- | --- |
| GOBP_CYTOKINE_MEDIATED SIGNALING_PATHWAY | MMP12, HCK, IL13RA2, CRLF2, MT3, CCL5, CXCR4, CXCL8, CCL3, HSPA1B, HSPA1A, LEP, CCL26, CCL19, IL36B |
| GOBP_REGULATION_OF_RESPONSE TO_EXTERNAL_STIMULUS | MAS1, MMP12, HCK, MT3, CCL5, CXCR4, ADORA2A, CXCL8, SERPINB2, MMP3, FGF18, CCL3, LEP, TNFSF14, CCL26, CCL19, MMP8, PDE2A, SEMA4D, NINJ1, SERPINF2, PTGES, TNFAIP6, TNFRSF1B, SIRPA |
| GOMF_CYTOKINE_RECEPTOR BINDING | CRLF2, CCL5, CXCL8, CCL3, TNFSF14, CCL26, CCL19, IL36B |
| KEGG_CHEMOKINE_SIGNALING PATHWAY | HCK, CCL5, CXCR4, CXCL8, CCL3, CCL26, CCL19 |
| GOBP_POSITIVE_REGULATION_OF IMMUNE_SYSTEM_PROCESS | MMP12, HCK, IL13RA2, CRLF2, CCL5, CXCL8, CCL3, CD247, LEP, TNFSF14, CCL19, IL36B, IGFBP2, BDKRB1, VNN1,MMP8, SIRPB1, GATA2 |
| GOBP_RESPONSE_TO_CYTOKINE | MMP12, HCK, IL13RA2, CRLF2, MT3, CCL5, CXCR4, CXCL8, CCL3, HSPA1B, HSPA1A, LEP, INA, CCL26, CCL19, IL36B, SLC2A4, PDE2A, GCH1, RBM15, TNFRSF1B, RAB20, SIRPA, FZD4 |
| GOBP_INFLAMMATORY_RESPONSE | MAS1, HCK, CCL5, CXCR4, ADORA2A, CXCL8, MMP3, CCL3, TREM1, LEP, CCL26, CCL19, IL36B, BDKRB1, VNN1, MMP8, PDE2A, NINJ1, SERPINF2, PTGES, TNFAIP6, TNFRSF1B, SIRPA |
| GOBP_DEFENSE_RESPONSE | MAS1, MMP12, HCK, CCL5, CXCR4, ADORA2A, CXCL8, MMP3, CCL3, ASIC4, TREM1, LEP, CCL26, CCL19, IL36B, PRG2, BDKRB1, VNN1, MMP8, PDE2A, GCH1, NINJ1, SERPINF2, PTGES, TNFAIP6, WFDC2, TNFRSF1B, RAB20, SIRPA |
| GOMF_EXTRACELLULAR_MATRIX STRUCTURAL_CONSTITUENT CONFERRING_TENSILE_STRENGTH | COL24A1, COL14A1, COL15A1, COL4A5, COL6A5, COL21A1, COL6A6 |
| GOCC_COLLAGEN_TRIMER | COL24A1, COLEC12, COL14A1, C1QTNF2, COL15A1, COL4A5, COL6A5, COL21A1, COL6A6 |
| GOCC_COLLAGEN_CONTAINING EXTRACELLULAR_MATRIX | POSTN, VCAN, EDIL3, FN1, ANXA2, COL24A1, ANGPT1, COL14A1, COL15A1, COL4A5, FGFR2, THSD4, COL6A5, ASPN, INHBE, COL21A1, SMOC2, COL6A6, OGN, F13A1, OMD |
| GOBP_CELL_CELL_ADHESION VIA_PLASMA_MEMBRANE ADHESION_MOLECULES | NEXN, PCDHA3, PCDHA10, PCDHAC1, PCDHA6, PCDHA7, PCDHA4, PCDHA11, PCDHA12, PCDHA8, PCDHA9, PCDHA1, PCDHA2, PCDHA5, PCDHA13, PCDHAC2, FLRT3 |

**Fig. S1A: A high concentration of talabostat induces epidermal atrophy in SEs in an ASC-dependent manner.** Quantification of IL-1β release and thickness of the epidermis of SEs upon treatment with talabostat (10 μM) for 3 days with daily medium change with fresh talabostat. Data are representative of at least 3 independent experiments. P values were calculated with two-way ANOVA analysis. (∗∗∗∗P < 0.0001, ∗∗∗P ≤ 0.001, ∗∗P ≤ 0.01, and ∗P ≤ 0.05, ns = not significant).

**S1B,C NLRP1 activation in SEs induces a pro-inflammatory signature.** SEs with control or ASC knockout HPKs (triplicates) were mock- or talabostat-treated (0.3 μM) for 3 days. Epidermis and dermis were separated and mRNA expression was characterized by NGS. **(B)** Heatmap of differentially expressed genes in epidermis and dermis of control mock-treated SEs demonstrating efficient separation of the skin layers and similar regulation of gene expression in the triplicates. **(C)** Leading edge genes (tables) are those that contribute most to the enrichment signal of a given gene set of pathways. Table indicating leading genes upon GSEA of NLRP1 activation in control SE and ASC knockout SE vs inflammatory pathways. The upper table is for epidermis and the lower for dermis.

TA: talabostat.

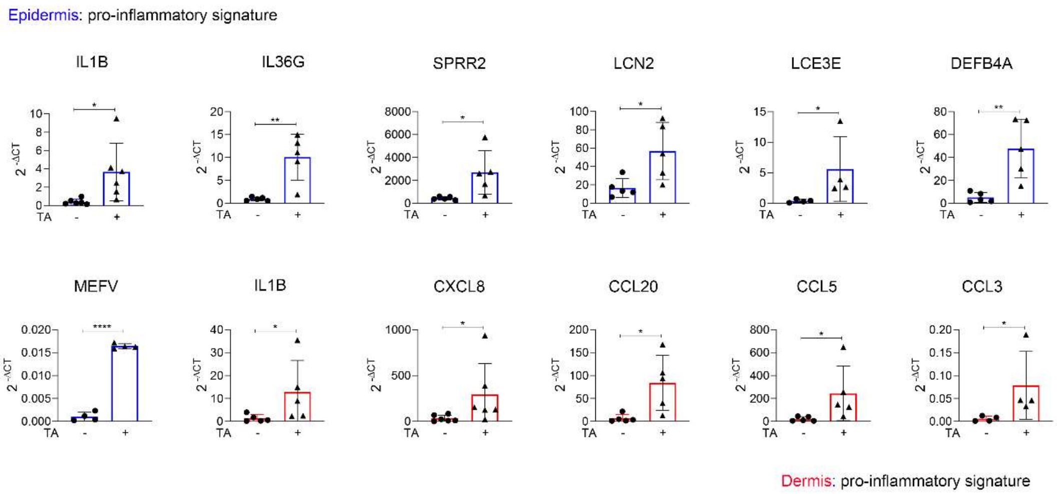


**Fig. S2: The NLRP1-dependent upregulation of pro-inflammatory genes is donor independent.**

SEs generated with HPKs from 5 different donors were mock-treated or treated with talabostat (0.3 μM) for 3 days. Expression of the indicated genes by keratinocytes (blue) or fibroblasts (red) was analyzed by qPCR and normalized to expression of the ratio between HPRT and RPL27. Data represent the average between 2 replicates of at least 4 different donors. P values were calculated with one-tailed unpaired t-test. (∗∗∗∗P < 0.0001, ∗∗∗P ≤ 0.001, ∗∗P ≤ 0.01, and ∗P ≤ 0.05, ns = not significant).

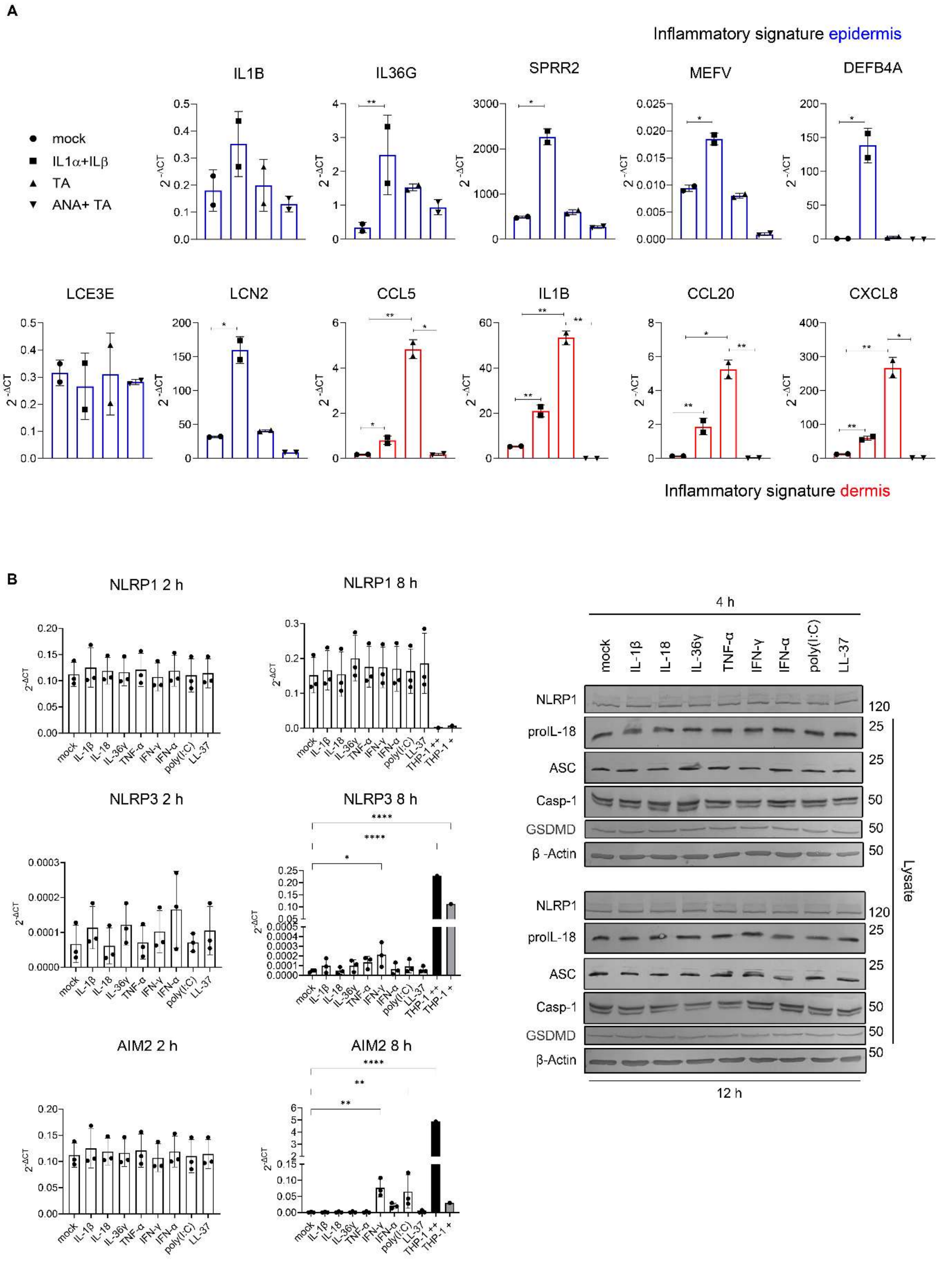
A
Inflammatory signature epidermis
IL1B
IL36G
SPRR2
MEFV
DEFB4A
mock
IL1α+ILβ
TA
ANA+ TA
2 -ΔCT
LCE3E
LCN2
CCL5
IL1B
CCL20
CXCL8
Inflammatory signature dermis
B
NLRP1 2 h
NLRP1 8 h
NLRP3 2 h
NLRP3 8 h
AIM2 2 h
AIM2 8 h
mock
IL-1β
IL-18
IL-36γ
TNF-α
IFN-γ
IFN-α
poly(I:C)
LL-37
THP-1 ++
THP-1 +
4 h
NLRP1
proIL-18
ASC
Casp-1
GSDMD
β -Actin
Lysate
12 h

**Fig. S3: IL-1 is the major effector of NLRP1 inflammasome activation.**

**(A)** SEs were mock-treated or with IL-1α/β (1 ng/ml each), talabostat (0.3 μM), or anakinra (10 μg/ml) plus talabostat, for 1 day. Expression of mRNA of representative genes of the epidermis (blue) and of the dermis (red) was determined by qPCR, relatively to HPRT expression. **(B)** Starved HPKs from 3 independent donors in monolayer were treated with IL-1β (10 ng/ml), IL-18 (20 ng/ml), IL-36γ (100 ng/ml), TNFα (10 ng/ml), IFN-γ (20 ng/ml), IFN-α (10 ng/ml), poly(I:C) (1 μg/ml), or LL-37 (1 μg/ml), and NLRP1, NLRP3 and AIM2 expression was determined at the RNA level 2 h and 8 h after stimulation (left) and at the protein level after 4 h and 12 h (right). THP-1 + (PMA 50 ng/ml) and THP-1 ++ (PMA 50 ng/ml, LPS 0.1 μg/ml) are used as positive control for NLRP3 and AIM2 RNA expression.

Data are represented by mean ± SD of **(A)** 2 replicates and are representative of 2 independent experiments or **(B)** 3 different donors, or a representative blot out of 3 is shown. P values were calculated with one-way ANOVA (∗∗∗∗$P < 0.0001$, ∗∗∗$P \leq 0.001$, ∗∗$P \leq 0.01$, and ∗$P \leq 0.05$, ns = not significant).

TA: talabostat; ANA: anakinra.

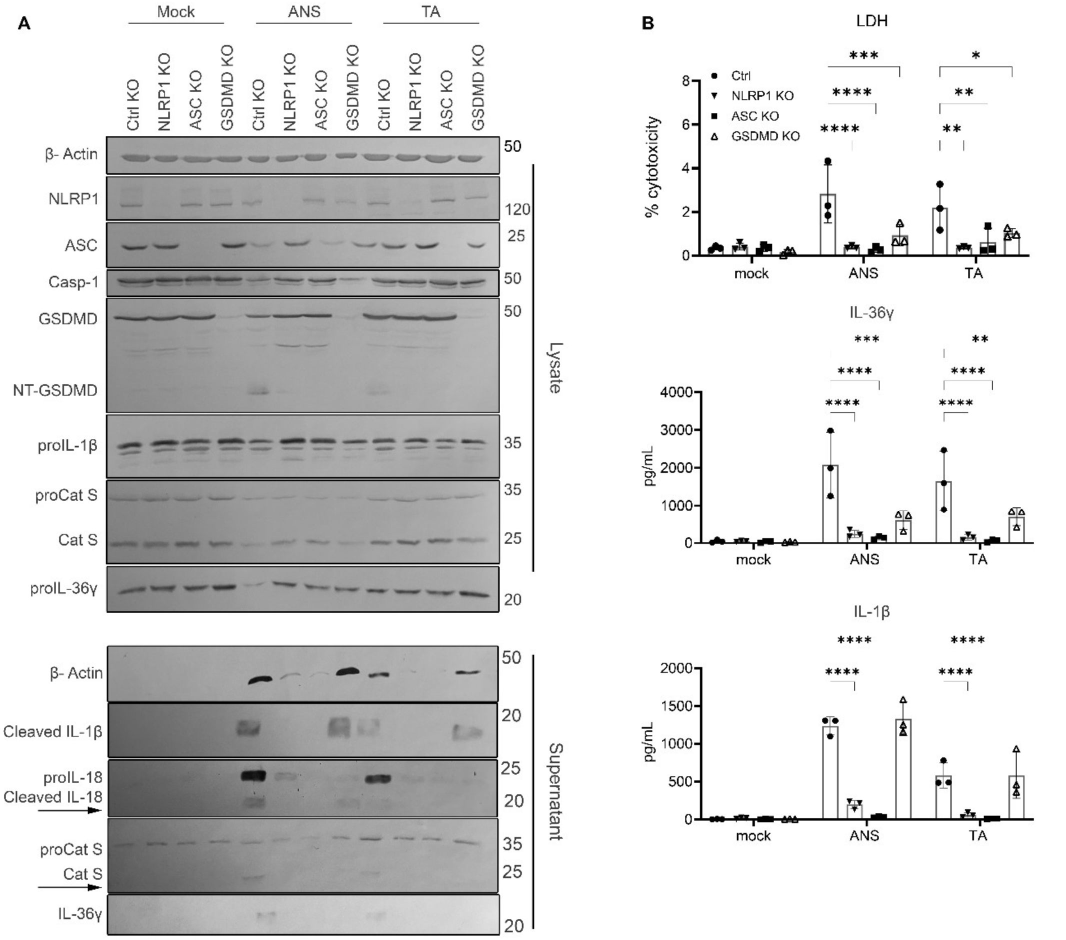


C

Leading edge genes
Ctrl SE TA Epidermis vs psoriatic epidermis

| | |
|---|---|
| BASAL2 UP | DEFB4A, CXCR4, LCN2, SPRR2A SPRR2E, CHI3L2, VNN1, IL36G, SPRR2D, KYNU, CHST2, RAC2, SLA, SERPINB4, S100A7A, CNFN, CD68, SLCO4A1, PI3,SERPINB3, ADIRF, OLFM1, LRG1, ALOX15B, DLX3, KRT75, CSTB |
| MITOTIC UP | DEFB4A, CXCR4, LCN2, SPRR2A, SPRR2E, CHI3L2, VNN1, IL36G,RHCG, CSTB, SPRR2D, KYNU, CHST2, RAC2, SLA, MUCL1, iL1B, SERPINB4, S100A7A, CNFN, CD68, PI3, SERPINB3, ADIRF, TGM1,OLFM1, LRG1, ALOX15B, DLX3, KRT75, |
| BASAL1 UP | DEFB4A, CXCR4, LCN2, SPRR2A,SPRR2E, CHI3L2, VNN1, IL36G, SPRR2D, KYNU, CHST2, SLA, IL1B, SERPINB4, S100A7A, CNFN, CD68, PI3, SERPINB3, ADIRF, TGM1, OLFM1, LRG1, ID3, ALOX15B, KRT75, CSTB |
| FOLLICULAR UP | DEFB4A, CXCR4, SPRR2A,SPRR2E, VNN1, IL36G, SPRR2D, RAC2, S100A7A, PI3 |
| GRANULAR UP | DEFB4A, SPRR2A, IL36G, RHCG, SPRR2D, SERPINB4, S100A7A, CD68, TGM1, LRG1 |
| CHANNEL UP | DEFB4A, SPRR2A, SPRR2E, CHI3L2, VNN1, IL36G, RHCG, SPRR2D, KYNU, SERPINB4, S100A7A, CNFN, PI3, SERPINB3, ADIRF |
| CHANNEL DOWN | HCFC1R1,CCND1,GLTP, PLIN2, CD24, SCCPDH, EFEMP1, SULF2, TNFRSF19, C12orf75, LURAP1L, WNT4, LAMB4, SYNPO2, RERG, KLK1 |
| BASAL1 DOWN | GLTP, PLIN2, TMEM40, ZNF165, SPTLC2, PDGFC, WNT3, EFEMP1, SULF2, KITLG, CA2, TNFRSF19, RASSF9, SOX6, BBOX1, SYNE1, LAMB4, SYNPO2, LRRN4CL,RERG |
| MITOTIC DOWN | BACE2, CCND1, GLTP, PLIN2, TMEM40, GRB7, SPTLC2, PDGFC, WNT3, SCCPDH, EFEMP1, SULF2, TCEA3, CABLES1, KITLG, CA2, TNFRSF19, RASSF9, HNMT, RASSF6, PTGS1, SOX6, BBOX1, SYNE1, LAMB4, SYNPO2 ,TNFRSF18, LRRN4CL,RERG |
| BASAL2 DOWN | WNT3, GPR155, SCCPDH, EFEMP1, SULF2, TCEA3, CABLES1, KITLG, CA2, TNFRSF19, RASSF9, HNMT, RASSF6, PTGS1, LURAP1L, SOX6, BBOX1, SYNE1, LAMB4, SYNPO2, TNFRSF18, MYLK, LRRN4CL, RERG, CLU, ASPN |

Leading edge genes
Ctrl SE TA Epidermis vs atopic dermatitis

| | |
|---|---|
| ATOPIC_DERM KERA_UP | CA2, TNFRSF19, LURAP1L, LAMB4, ASPN |

**Fig. S4A,B: Active cathepsin S is secreted upon NLRP1 inflammasome activation.**

**(A)** Polyclonal knockout HPKs from 3 independent donors in 2D monolayer were primed with IL-1α (10 ng/ml) and IFN-γ (20 ng/ml) and mock-treated or with anisomycin (1 μM, 5 h), or talabostat (10 μM, 12 h). Protein expression and release was determined by **(A)** western blot or **(B)** ELISA**. (B)** Cell death was measured by LDH release**.** Data are represented by mean ± SD of 3 independent donors or a representative blot out of 3 experiments is shown. P values were calculated by two-way ANOVA (∗∗∗∗$P < 0.0001$, ∗∗∗$P \leq 0.001$, ∗∗$P \leq 0.01$, and ∗$P \leq 0.05$, ns = not significant).

**S4C: Leading genes of regulated pathways upon GSEA of epidermis of talabostat-treated control SEs vs epidermis in psoriasis or atopic dermatitis.**

ANS: anisomycin; TA: talabostat.

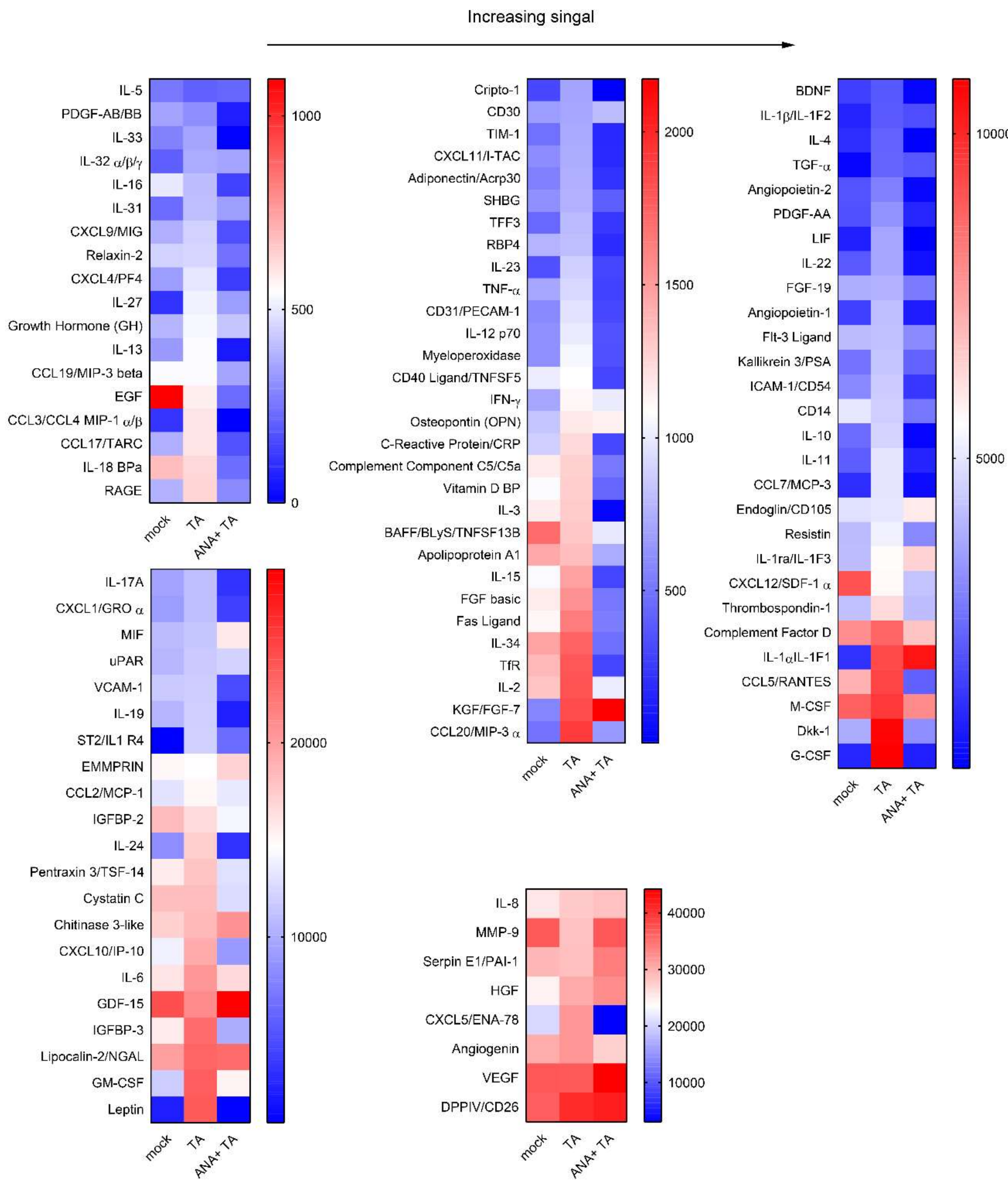


**Fig. S5: NLRP1 activation regulates release of several inflammatory cytokines.**

SEs with wild type HPKs were mock-treated or treated with talabostat (0.3 µM) or with anakinra (10 µg/ml) plus talabostat, for 3 days. Release of the indicated proteins was determined by cytokine array in a semi-quantitative manner.
